# ATP synthase interactome analysis identifies Mco10 – a new modulator of permeability transition pore in yeast

**DOI:** 10.1101/2022.07.02.498517

**Authors:** Chiranjit Panja, Aneta Wiesyk, Katarzyna Niedźwiecka, Emilia Baranowska, Roza Kucharczyk

## Abstract

In *S. cerevisiae*, the uncharacterized protein Mco10 (**M**itochondrial **c**lass **o**ne protein of **10** kDa) was previously found to be associated with mitochondrial ATP synthase and referred to as a new ‘subunit <u>*l*</u>’. However, recent cryo-EM structures of *S. cerevisiae* ATP synthase could not ascertain Mco10 as a structural subunit of the enzyme, either monomers or dimers, making questionable its role as a structural subunit. The N-terminal part of Mco10 is very similar to Atp19 (subunit *k*) of ATP synthase. The subunit *k*/Atp19, along with the subunits *g*/Atp20 and *e*/Atp21 plays a major role in stabilization of the ATP synthase dimers. In our effort to confidently define the small protein interactome of ATP synthase we similarly found Mco10 associated with ATP synthase of *S. cerevisiae*. We herein investigated the impact of Mco10 on ATP synthase functioning. Biochemical analysis revealed in spite of similarity in sequence and evolutionary lineage, that Mco10 and Atp19 differ significantly in function. This is the first work to show Mco10 is an auxiliary ATP synthase subunit that only functions in permeability transition.

## Introduction

Mitochondria, the dynamic organelles of endosymbiotic origin, are the source of the cellular energy in the form of ATP necessary for life and at the same time they are the origin of intrinsic signals directing the cell into programmed death pathway (Tait & Green, 2013; Vakifahmetoglu-Norberg et al., 2017). Apart from ATP synthesis by the ATP synthase complex in the process of oxidative phosphorylation, OXPHOS (Saraste, 1999), mitochondria is the hub for synthesis of metabolites, Fe-S clusters and essential amino acids (Spinelli & Haigis, 2018). The *Saccharomyces cerevisiae* as a model have provided fundamental insights into mitochondrial biology due to its ability to survive under fermenting conditions when OXPHOS is inactivated due to mutations in the structural and regulatory proteins of this system (Chen & Clark-Walker, 2000).

The ATP synthase monomer complex of *S. cerevisiae* is composed of 17 known subunits. It is organized into a membrane-embedded F_O_ domain and membrane-extrinsic F_1_ catalytic domain connected by two stalks: central and peripheral stalk. The F_O_ domain is built from the ring of ten subunits *c*/Atp9, subunit *a*/Atp6 and the small subunits Atp8, Atp18/*i*, Atp17/*f*, and membrane part of subunit Atp4/*b*. The matrix part of subunit *b* together with subunits Atp14/*h*, Atp7/*d*, Atp5/*OSCP* forms the external stalk connecting the F_O_ with the top of the F1. The hydrophilic F_1_ domain built from the hexamer of subunits *α*_3_*β*_3_ and the central stalk built from subunits Atp3/γ, Atp16/δ and Atp15/ε, are connected to the *c*-ring by subunit γ (Guo et al., 2017). The ATP synthase monomers form dimers connected by subunits Atp6/*a* interaction (Velours et al., 2011) and stabilized by the supernumerary subunits (subunits that are not essential for the catalytic activity) Atp21*/e*, Atp20/g and Atp19/*k* (Arnold et al., 1998; Brunner et al., 2002; Paumard et al., 2002). Very little is known about the role of Atp19. It is the homolog of mammalian subunit *k*/DAPIT, knock- out of which reduces ATP synthesis and destabilize the dimers (Arnold et al., 1998). Similarly, deletion of Atp19 reduces dimers formation in *S. cerevisiae* (Wagner et al., 2010). Mutation in the gene encoding subunit *k* has been reported to result in Leigh syndrome in humans due to the reduced dimer formation and impaired ATP synthesis (Barca et al., 2018).

Most of the ATP synthase subunits are small proteins in the range 5-20 kDa. In addition to the structural subunits *c, 8, d, h, i, f, δ* and *ε*, this group includes hydrophobic super numeracy subunits *e, g, k*. Also, most small regulatory proteins known to interact with ATP synthase and possibly modulates its function are within this molecular weight range (Ebanks et al., 2022). However, identification of these proteins by mass spectrometry and western blot remains difficult due to their very low abundance, poor fragmentation and hydrophobic nature (Ahrens et al., 2022). In this work, we analyzed extensively proteins in this molecular weight range that we defined as ‘small protein interactome of ≤ 20 kDa’ of ATP synthase to well understand these mini/micro structural and regulatory proteins that bind to ATP synthase in *S. cerevisiae*. In an effort to differentiate how this interactome varies when ATP synthase undergoes dimerization, we further analyzed the subunit composition and proteins that remains associated with the monomers and dimers of ATP synthase in *S. cerevisiae* and also compared the interactome with human ATP synthase. This work focuses on characterizing the function of one protein Mco10 (**M**itochondrial **c**lass **o**ne **p**rotein of **10** kDa) encoded by the *YOR020W-A* gene in *S. cerevisiae* identified in our interactome data to be associated with the ATP synthase complex.

Mco10 and its homolog in *P. angusta* were also previously identified with ATP synthase in a pull-down experiment and was also named as a new subunit ‘*l*’ (Liu et al., 2015; Morgenstern et al., 2017). The N-terminal part of this protein is very similar in sequence to subunit Atp19/*k*, homologs of which were also present in those species. Interestingly, Mco10 and its homolog were not present in subsequent cryo-EM structures of ATP synthases from *S. cerevisiae* and *P. angusta*, thus the demonstration that Mco10 to be a bona fide subunit of the ATP synthase complex is missing (Guo et al., 2017; Srivastava et al., 2018; Vinothkumar et al., 2016). In this work, we present evidence that Mco10 can indeed be a true subunit of ATP synthase in *S. cerevisiae*. Phylogenetic analysis of Mco10 and Atp19 in fungal species further showed that many related yeast species maintain the two proteins and Mco10 is evolutionarily more ancient. However, Mco10 and Atp19 functions are significantly different. Mco10 has a role in modulating the permeability transition pore (PTP, also called the mitochondrial megachannel) and calcium homeostasis in *S. cerevisiae*, unlike Atp19, which we found does not modulates PTP.

In the last decade, several groups demonstrated the involvement of ATP synthase in PTP formation. However, despite extensive research in the field, the exact composition and molecular mechanism of the PTP still remain a mystery to be solved (Bernardi, 2020; Bernardi et al., 2021). The PTP is defined as a Ca^2+^-activated channel of the inner mitochondrial membrane which mediates the permeability to solutes up to 1.5 kDa (Azzolin et al., 2010; Haworth & Hunter, 1979). It was demonstrated that the PTP resides in the *c*-ring and is gated by the F_1_ dissociation (Alavian et al., 2014; Mnatsakanyan et al., 2019). The ATP synthase dimer junction may also be involved (Carraro et al., 2014; Niedzwiecka et al., 2018). The Ca^2+^ ions binding to the β subunit, the OSCP subunit involvement in reactive oxygen species modulation of PTP was demonstrated. The adenine nucleotide transporter (ANT) was also found to contribute to the pore opening (Carraro et al., 2020; Giorgio et al., 2017; Neginskaya et al., 2019). The primary consequence of PTP activation, when it opens for a long time, is the inner mitochondrial membrane depolarization followed by matrix swelling and rupture of the outer membrane. This process releases the pro-apoptotic factors and is being increasingly appreciated to be at the center of cellular death cascade (Bauer & Murphy, 2020; Ichas & Mazat, 1998; Lemasters et al., 2009; Mnatsakanyan et al., 2022). Search of a pharmacological targets to inhibit PTP as a cure in several neuro-muscular diseases is a very active area of research (Antonucci et al., 2020; Boyenle et al., 2022). The fact that an unknown small protein like Mco10 can modulate Ca^2+^ homeostasis and PTP through its association to a well-studied enzyme like ATP synthase points to the important role these small proteins play in physiological pathways (He et al., 2018).

## Results

### Small protein interactome of ATP synthase

The full ATP synthase complex was pulled down by HA-6xHis-tagged Atp6 bound to Ni-NTA agarose. The proteins were then eluted from the beads, separated in the 15 % SDS PAGE gel and visualized by silver staining. In another approach, the monomers and dimers of ATP synthase were extracted from isolated mitochondria by 2% digitonin, separated in 3-12% BN-PAGE, and then the monomers and dimers were resolved in a 2^nd^ dimension 16 % SDS PAGE **(Fig 1A)**. To compare the yeast interactome with human, the monomers and dimers of ATP synthase from mitochondria isolated from HEK293T were extracted with similar digitonin concentration as from yeast and similarly resolved in 2D- BN-SDS PAGE. Gel lane fragments in the 5-20 kDa range were submitted for mass spectrometry identification as defined in the methods section **(Appendix Fig S1)**.

**Figure 1.**
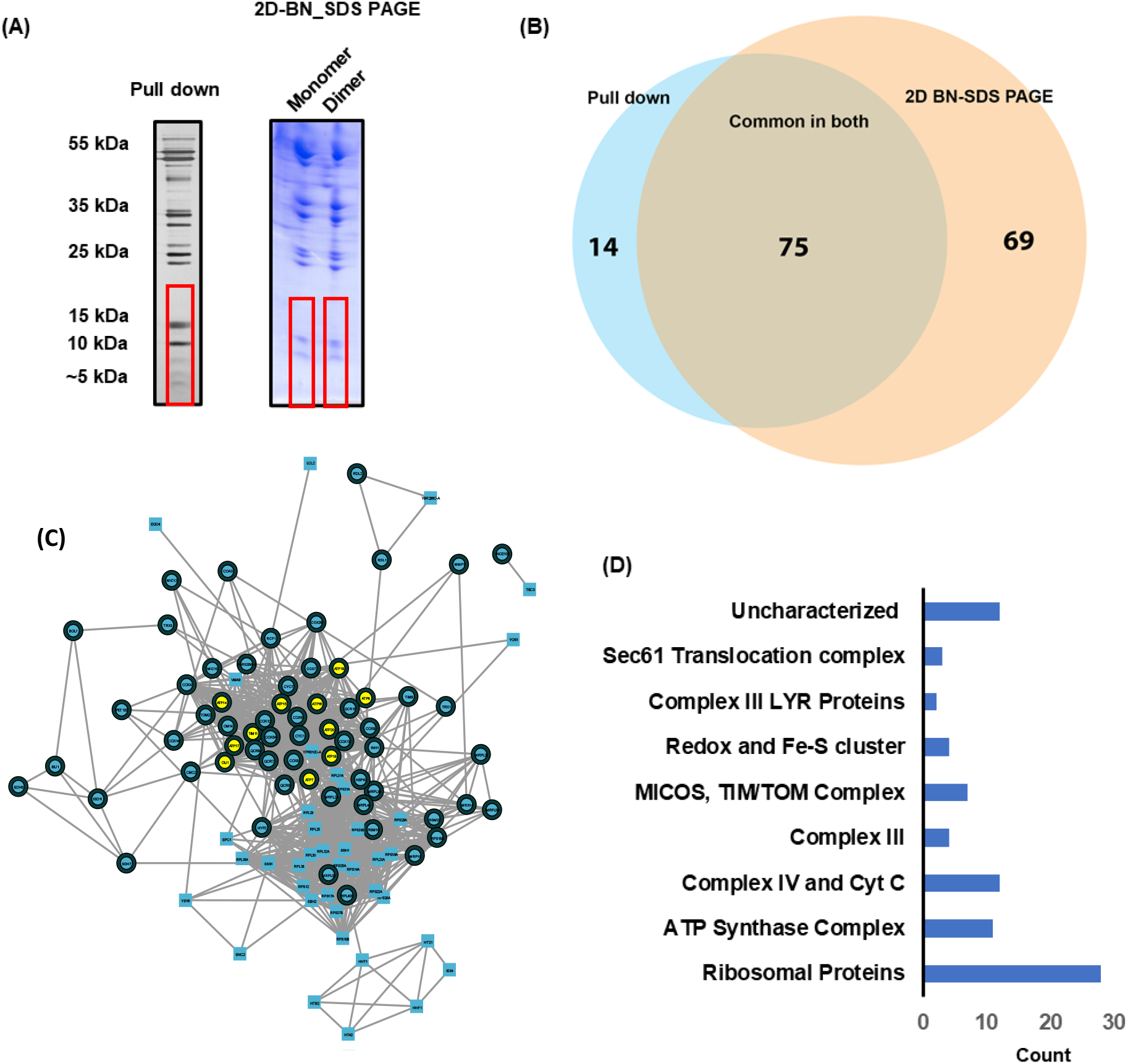
Small molecule interactome of ATP synthase of S. *cerevisiae*. **A**. Pulldown of ATP synthase complex by HA-6His-tagged AtpS or the monomers and dimers of ATP synthase extracted by digitonin and separated in twodimensional BN-SDS-PAGE gel and visualized by silver stain or Coomassie Gel pieces from region marked in red were cut-off for LC/MS analysis. The experiments were performed many times and representative gels are shown. **B**. Venn diagram of the < 20 kDa proteins associated with ATP synthase identified in the interactome analysis from the two different approaches. **C**. High confidence ATP synthase interactors identified by MS were connected into a network using the STRING database and visualized using the STRING 106 plugin in Cytoscape. Proteins marked in circle are known proteins that localizes to the mitochondria. Proteins marked in yellow circle are the known ATP synthase subunits identified in the analysis. **D**. Classification of the identified proteins into known complexes and processes.

From the analysis of *S. cerevisiae* samples, a total of 89 proteins from pull down experiment and 144 proteins from 2D-BN-SDS PAGE got enriched according to the set criteria outlined in methods. Among them, 75 proteins were commonly identified in both the experiments **(Fig 1B)**. All the eleven subunits of ATP synthase with mol. wt ≤ 20 kDa also got identified including the ATPase activity inhibitor protein Inh1. However, Atp8 were only identified in pull down and not detected from 2D-BN-PAGE samples. The result also showed that the three known supernumerary subunits Atp19/*k*, Atp20/*g* and Atp21/*e* were present in both the monomer and the dimer. The yeast interactome was also highly enriched with several ribosomal proteins which were more abundant in 2D-BN-SDS PAGE. Also, we found several subunits of Complex IV including Cyt C and four members of Complex III to remain associated with ATP synthase in both approaches showing they form sub-complexes with ATP synthase. In addition, members of the Sec61 translocation complex, the chaperonin Hsp10 and member of the MICOS component Mic10 were identified in both the monomers and the dimers. Mic10 was previously shown to interact with the dimeric F_1_Fo-ATP synthase (Rampelt et al., 2017). Three uncharacterized proteins Yor020W-A (Mco10), Ypr010C-A (Min8), Yir021W-A, were enriched with ATP synthase in our study showing they can potentially modulate ATP synthase in yet unknown way **(Fig 1C and 1D, Table EV1)**.

The human ATP synthase interactome of small proteins identified 136 proteins among which 16 proteins were only identified in the monomers and 21 proteins got identified only in the dimers **(Fig EF1 and Table EV2)**. We identified all ten subunits of the ATP synthase complex and the ATPase activity inhibitor protein within this defined mol. wt range. However, subunits *g, c* and the coupling factor 6 were identified only in the dimer sample. Several members of Complex IV and Complex I were also highly enriched. Homologs of many proteins outside of the OXPHOS complexes, identified in yeast ATP synthase interactome, were also present in the human ATP synthase interactome, validating the yeast data. Examples include members of the Sec61 translocation complex, LYR motif containing biogenesis and assembly factors, thioredoxin and chaperonin Hsp10. Interestingly, the TIM/TOM complex was enriched in the human but got poorly represented in the yeast ATP synthase interactome. Another highly enriched cluster in the human interactome were members of the S100 Calcium binding complex, which we identified for the first time to interact with ATP synthase.

**Figure EF1.**
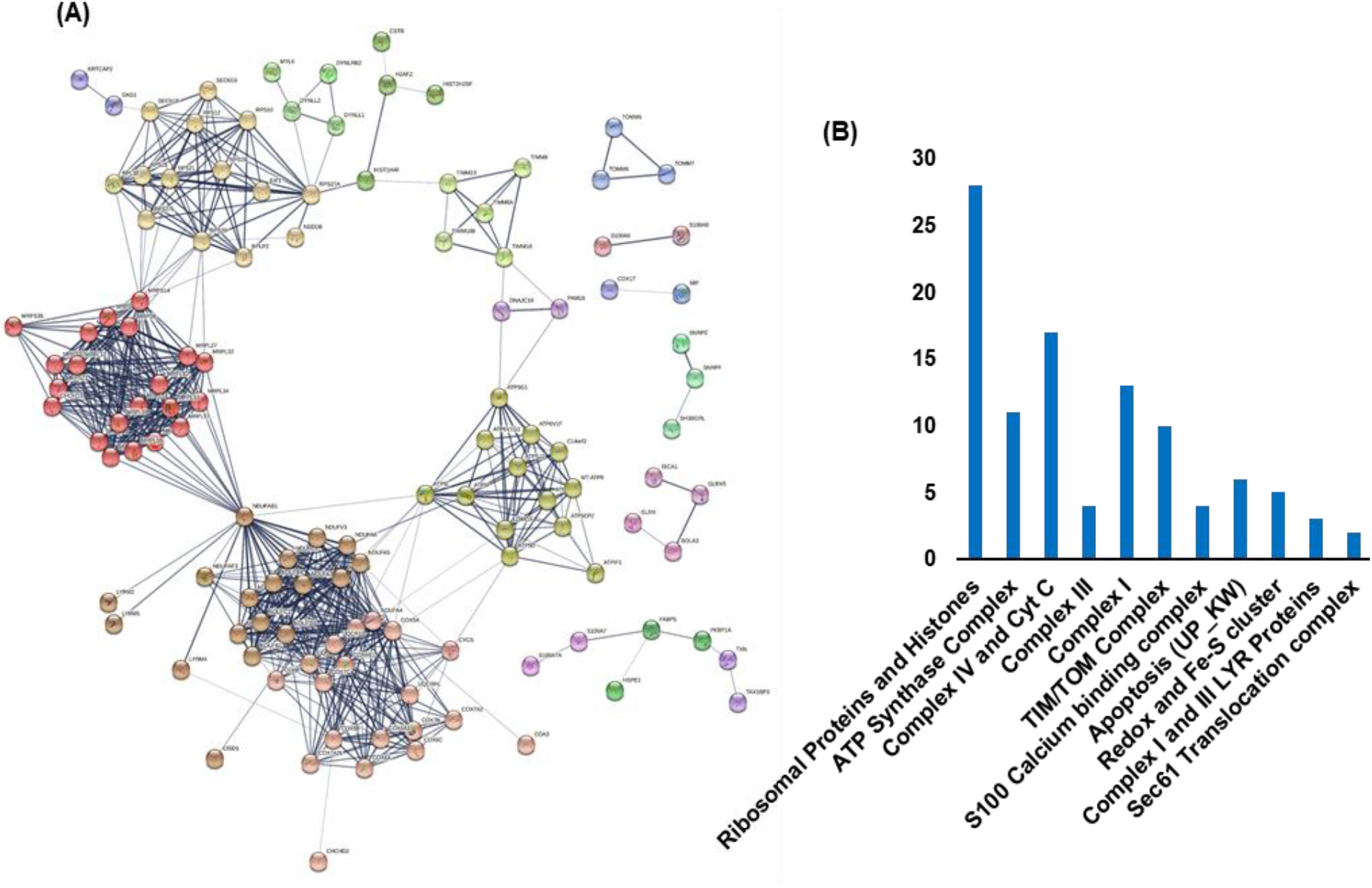
Human ATP synthase interactome. Mitochondria were isolated from HEK293T. The monomers and dimers of ATP synthase were extracted by digitonin and separated in two-dimensional BN-SDS-PAGE gel and visualized by Coomassie. Gel pieces from ≤ 20 kDa region from monomer and dimer were cut-off and analyzed by LC/MS analysis. Interactome was analyzed using the STRING database and categorized manually or by Gene Ontology terms.

### Yor020W-A (Mco10) co-purifies with the ATP synthase complex

From the pull down and 2D-BN-SDS PAGE analysis of the monomers and dimers of *S. cerevisiae* ATP synthase, the uncharacterized protein Yor020W-A (Mco10) was among the highly enriched and identified with more than 50 peptides and greater than 50% sequence coverage **(Appendix Fig S2)**. However, it was only identified in the monomer in two replicates. As Mco10 was previously identified in a pull down of ATP synthase and hypothesized as its potential subunit (Liu et al., 2015) we set to characterize the Mco10 role in ATP synthase function.

### Phylogenic analysis of Mco10 and Atp19 based on sequence similarity

Detailed comparison of the sequences of fungal subunits *k* and *l* showed that these proteins indeed have high similarity within their N-terminal part as noted previously (Liu et al., 2015). *S. cerevisiae* Mco10 and Atp19 share around thirty percent sequence similarity, mainly within the N-terminal region **(Fig 2A)**. The serine rich hydrophilic middle region is characteristic for Mco10 and is absent in Atp19 **(Fig 2B)**. The structures of Mco10 and Atp19 were analyzed using the available structures in AlphaFold2 database **(Fig 2C)**. The structures of related proteins in *Candida albicans* and *Pichia angusta* were also analyzed using predicted models available in AlphaFold2 database **(Fig EF2)**. Mco10 and Atp19 (and the related homologs) mainly consists of two helices with a disordered middle region. The N-terminal helical region that is similar in sequence in both proteins align perfectly (by PyMOL software). Although the exact spatial positions of the C-terminal helices remain difficult to predict due to the poor accuracy of the middle region, but analysis of AlphaFold’s expected position error PAE plot does give a fair confidence on their orientation in the predicted models **(Fig 2D)**. We performed an extensive phylogenetic analysis of the subunits *k* and *l* genes sequences and their related homologs that are present or are hypothesized to be present in the published genomes of fungi. We found that not only *S. cerevisiae* and *P. angusta*, but many fungi in the subphylum *Saccharomycotina* (Phylum *Ascomycota*) have homologs of both genes. But if we diverged out to other subphylum *Pezizomycotina* or *Taphrinomycotina* of the *Ascomycota*, or even in the *Basidiomycota*, we found mainly a single gene encoding subunit *k* or *l*. A phylogenetic tree of subunits *k* and *l* from all subphylum of *Ascomycota*, including *Basidiomycota* showed that both subunits originated from a common ancestor and subunit *l* is the more ancient. However, given the time scale these proteins have diverged, it is very likely that subunits *k* and *l* gained independent additional features with respect to ATP synthase structure and function **(Fig 2E, Appendix Fig S3, Table EV3)**.

**Figure 2.**
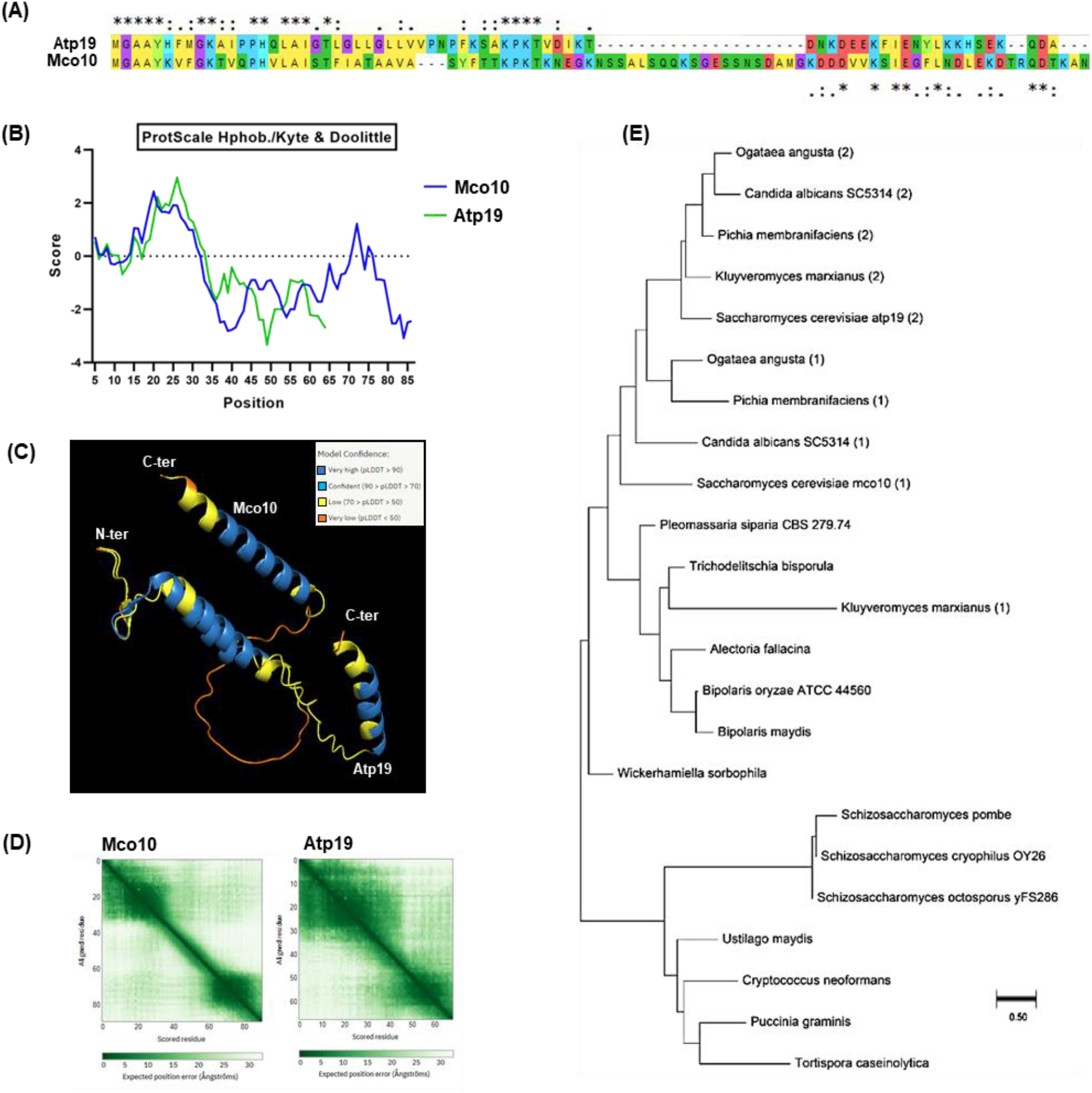
Mco10 and Atp19 phylogenetic and structural analysis. **A**. Alignment of Mco10 and Atp19 sequences by Clustal Omega. **B**. Hydropathy plots of Mco10 and Atp19 by Kate and Doolittle method using ProtScale on the ExPASy Server, **C**. Structures of Mco10 and Atp19 as predicted in the model from AlphaFold2 database. The color code represents the model confidence as determined by the Al algorithm of AlphaFold2. The alignment was done using PyMol molecular visualization software. **D**. AlphaFold’s expected position error PAE plot of Mco10 and Atp19. **E**. Phylogenetic tree analysis of Mco10, Atp19 and related homologs in fungi. The tree was constructed using Maximum likelihood method with representative species from all subphylum of Ascomycota and representative examples of Basidiomycota. (1) represents Mco10 related homolog and (2) represents Atp19 related homolog when there were two distinct proteins present in the aenomeofthe oraanism.

**Figure EF2.**
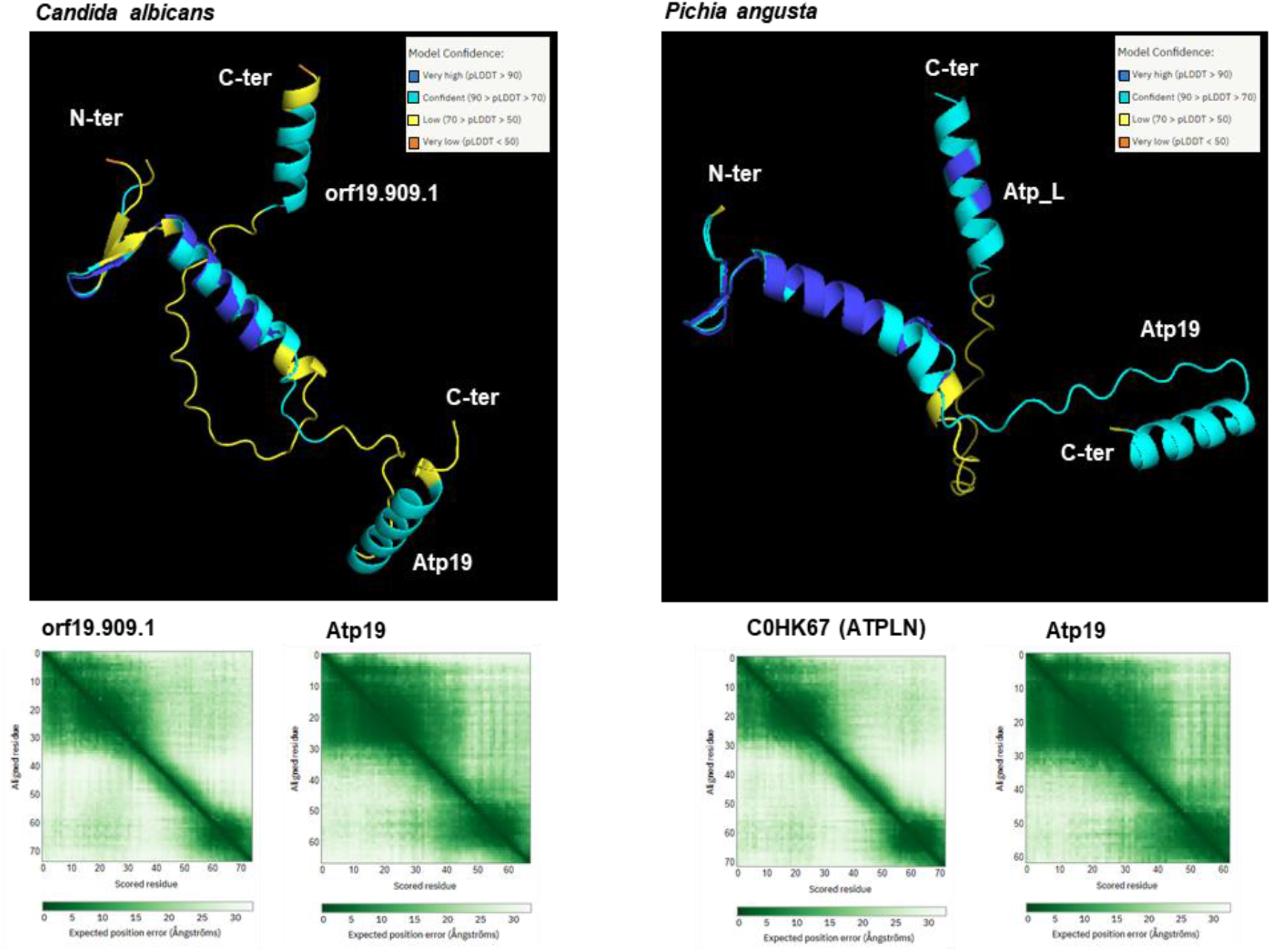
Superposition of subunits *k* and *I* from *Candida albicans* and *Pichia angusta*. The AlphaFold2 predicted structure of Atp19 (subunit *k)* and orf19.909.1 (subunit /) of *Candida albicans were* aligned using PyMol software. Color code represents the model confidence and PAE plots of the models are shown below. Homologous proteins from *Pichia angusta* were similarly modelled and visualized in PyMOL.

### ATP synthase activity and mitochondrial membrane potential

To check if Mco10 is indeed attached to ATP synthase we first checked the respiratory growth of *Δmco10, Δatp19* and the double deletion *Δatp19Δmco10* mutants. As shown on Fig 3A, the respiratory growth of these mutants is not affected at 28°C or even at 36°C, the conditions of moderate heat stress for yeast. Then we supplemented the respiratory medium with suboptimal concentration of oligomycin, an ATP synthase inhibitor, which does not affect growth of wild type strain. In these conditions deletion of Atp19 significantly reduces the growth at 1 µg/ml oligomycin at both growth temperatures. Deletion of Mco10 does not affect the respiratory growth under these conditions. Moreover, *Δmco10* was more resistant to oligomycin at 36°C and its deletion in *Δatp19* background, i.e., in the double mutant, also rescued the growth at 28°C to some extent **(Fig 3A)**.

**Figure 3.**
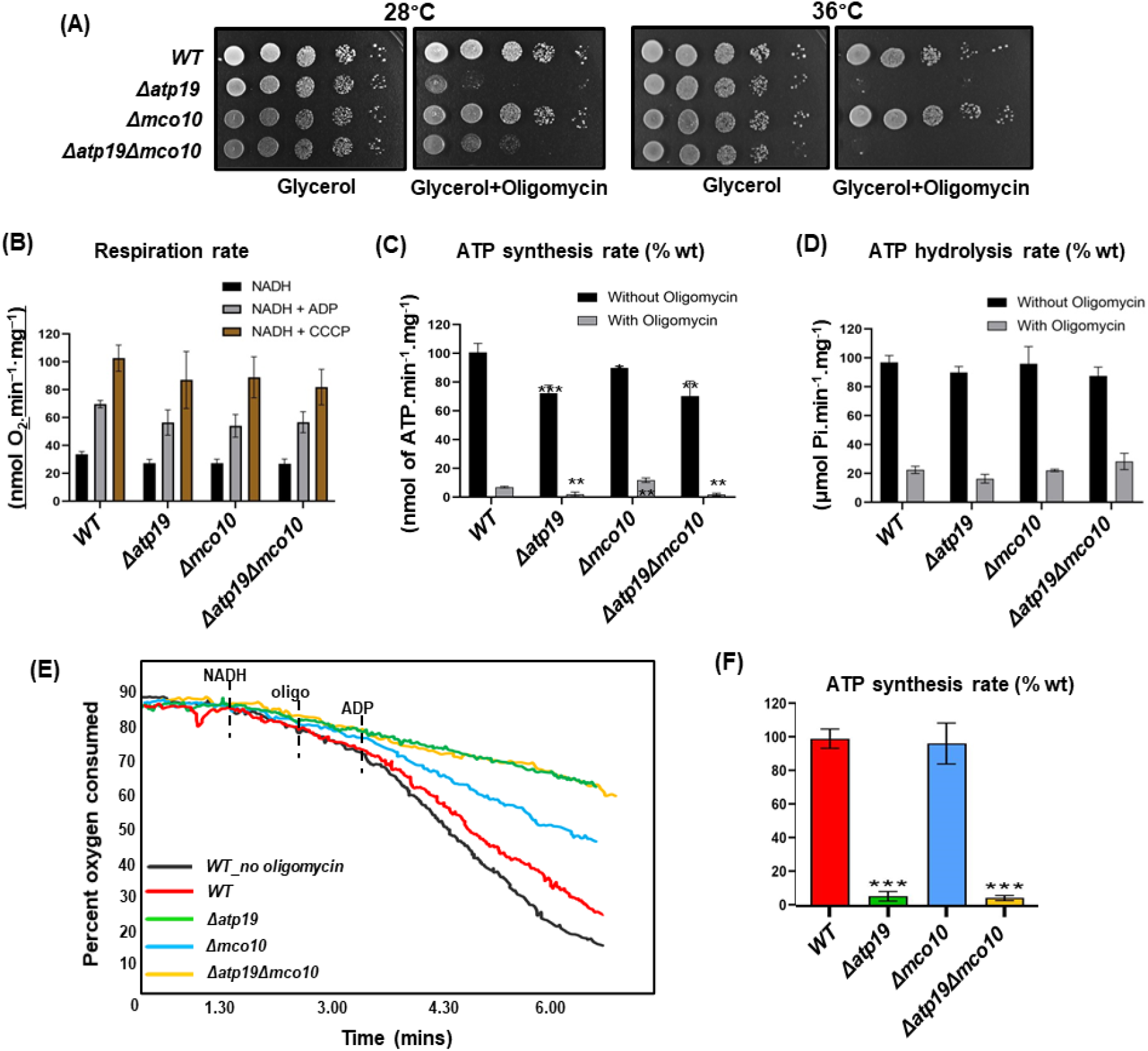
Respiratory growth, oxygen consumption rates, ATP synthase activities in *ΔmcolO, Δatp19* and *Δatp19Δmco10* mutants. **A**. Respiratory growth phenotypes. Cells from the indicated strains grown in glucose precultures were serially diluted and spotted on rich glucose or glycerol plates with or without oligomycin (1 pg/ml) and incubated at 28 or 36 °C. The glycerol plates were photographed after three days while the glycerol + oligomycin plates after four days of incubation. **B, C, D**. Oxygen consumption rates, ATP synthesis and hydrolysis activities Measurements were performed on freshly isolated mitochondria. The maximal oxygen consumption, at state 4 and state 3 are quantified when NADH, ADP and CCCP were added The respiration rate is expressed in percentage of the uncoupled respiration of wild type mitochondria. ATP synthesis and hydrolysis activities (darker rectangles) are expressed in percentage with respect to wild type mitochondria, whereas the activities in the presence of oligomycin (lighter rectangles) are expressed as the percentage of corresponding activities without oligomycin. **E**. The oxygen consumption rate of isolated mitochondria measured in the presence of 0.5 pg/ml oligomycin before addition of ADP **F**. ATP synthesis measured in the conditions of experiment shown in panel E, represented as the percentage of the ATP production in control mitochondria. Statistical significance is indicated by * p≤0.05, ** p≤0.005, *** p≤0.001.

We consequently asked whether lack of Mco10 affects the OXPHOS and ATP synthase activities in wild type and in *Δatp19* background. The oxygen consumption was measured in isolated mitochondria with NADH as an electron donor, alone (basal or state 4 respiration), and after successive addition of ADP (state 3, phosphorylating conditions) or CCCP (uncoupled, maximal respiration). At 28°C, *Δmco10, Δatp19* or the double deletion *Δatp19Δmco10* mutants does not significantly affect oxygen consumption rates of mitochondria in all above conditions **(Fig 3B)**. We next measured the rate of ATP synthesis by ATP synthase at the state 3 (in the presence and absence of 2.5 µg/ml concentration of oligomycin that totally blocks ATP synthase in wild type mitochondria). The ATP synthesis rates were decreased by 25% in *Δatp19* but not very significantly in *Δmco10*. However, in presence of oligomycin, which completely blocked ATP synthase in *Δatp19*, the rate was twice higher in *Δmco10* in comparison to wild type mitochondria **(Fig 3C)**. We assessed the functioning of ATP synthase in the reverse mode by measuring the rate of ATP hydrolysis in mitochondria not osmotically protected, so as to relax ATP synthase from any membrane potential that would limit its ATP hydrolytic activity, at pH 8.4 to avoid binding to F_1_ of its natural inhibitor IF1 (Venard et al., 2003). To assess the ATPase activity of ATP synthase the measurements were performed in the presence of oligomycin, which blocks rotation of *c*-ring and hence F_1_-mediated ATP hydrolysis. In wild type mitochondria the ATPase activity is inhibited by 80–90%. The ATPase activity in the mutants were not significantly affected, which is not surprising owing to the capacity of ATP synthase to assemble properly in absence of these subunits **(Fig 3D)**.

We then determined the rate of oxygen consumption and ATP production with a concentration of oligomycin at 0.5 µg/ml that inhibits the oxygen consumption by around 20% in wild type mitochondria isolated from strains grown at 28°C (**Fig 3E**). It should be noted, that at higher concentration, oligomycin can totally block ATP synthase in the wild type mitochondria. Under this suboptimal dose, *Δmco10* mitochondria retained ATP synthase function, albeit to a lower extent (50%) comparing to control mitochondria. But this concentration of oligomycin was sufficient to block ATP synthase in *Δatp19* and *Δatp19Δmco10* double mutant entirely **(Fig 3F)**.

We further investigated ATP synthase functionality in the mutant strains by membrane potential (ΔΨ) measurements, using Rhodamine 123, a cationic dye whose fluorescence is quenched when ΔΨ increases. For evaluating ΔΨ changes upon respiratory chain and ATP synthase activity, the mitochondrial membrane was first energized by feeding the respiratory chain with electrons from ethanol. Then ADP was added what results in a transient ΔΨ drop due to proton reentry which is reestablished within one minute. Then the ΔΨ is collapsed by addition of the complex IV inhibitor KCN but then ATP synthase consumes ATP produced during previous step of the experiment and pumps protons reestablishing partially the membrane potential **(Fig 4)**. The ΔΨ variations during experiment were not affected in *Δmco10* and *Δatp19* but the time of ΔΨ recovery after ADP addition in the double *Δatp19Δmco10* mutant was longer. This experiment confirms a defect of about 10% in oxygen consumption and 10-20% in ATP synthesis/hydrolysis activities observed in above experiments in mitochondria lacking both Mco10 and Atp19 subunits.

**Figure 4.**
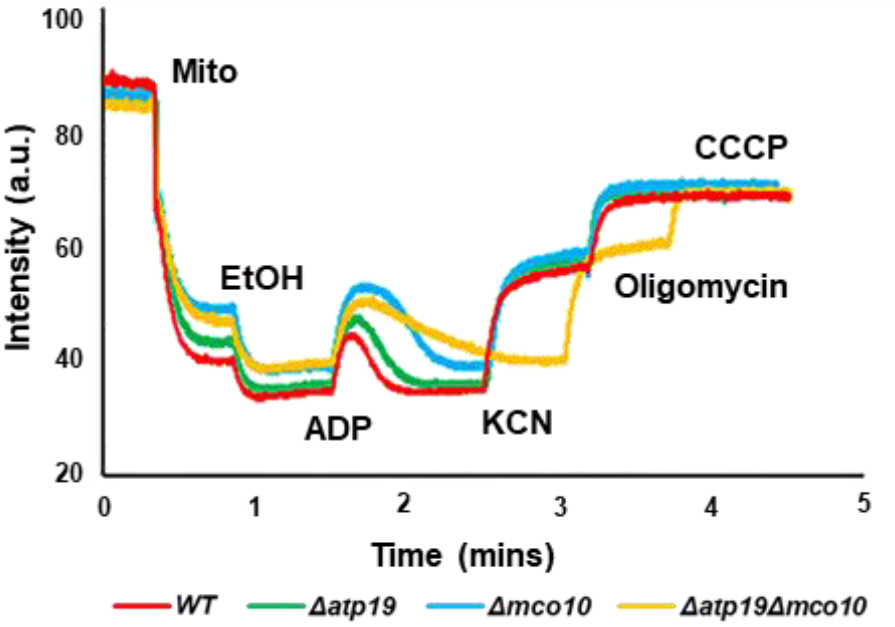
Variations in mitochondrial membrane potential in isolated mitochondria from strains grown at 28 °C measured under osmotic protection. The additions were 0.5pg/mL Rhodamine 123, 150pg/mL of mitochondrial proteins (Mito), 10pL ethanol (EtOH), 75 pM ADP, 2mM potassium cyanide (KCN), 4 pg/mL oligomycin (oligo), and 4 pM carbonyl cyanide-m-chlorophenyl hydrazone (CCCP).

### Assembly/stability of ATP synthase in *Δmco10* mutant

In order to determine the Mco10 and Atp19 impact on ATP synthase complexes we extracted them from mitochondria prepared from *Δatp19, Δmco10* and the double deletion *Δatp19Δmco10* mutants and separated on 3-12 % Blue Native PAGE, followed by Western blotting. Then 500 µg of mitochondria were incubated in extraction buffer containing 2% of digitonin for time period varying from 20 to 60 mins. With higher digitonin concentration and/or longer incubation time the stability of dimers decreases (our observation based on the experiments performed on mitochondria from W303-1B strain background routinely used in our laboratory). However, in the BY4741 wildtype strain mitochondria, the ATP synthase dimeric and monomeric complexes were stable after 60 minutes extraction time independently on the *Δmco10, Δatp19* single and the double *Δatp19Δmco10* deletion **(Fig 5A)**. Although the BN-PAGE technique is a semi-quantitative, from the ratio of dimers to monomers we conclude that much less dimers is extracted from mitochondria lacking Atp19 but not Mco10 at each extraction time. These results confirm the Atp19 role in stabilization of the dimers, as proposed by other researchers (Wagner et al., 2010). In contrast, when Mco10 is absent more ATP synthase complexes, i.e., both the monomers and the dimers were extracted after 20 and 30 minutes comparing to the wild type enzyme with 2% digitonin **(Fig 5A and Fig EF3)**. From this we conclude that ATP synthase complexes are extracted more easily in *Δmco10* mitochondria. Interestingly, more oligomers were also extracted from the mitochondria of the double mutant at 30 min extraction time **(Fig 5A)**. The steady state level of Atp6 and Atp2 subunits was also not changed in mitochondria in *Δmco10, Δatp19* single and the double mutant **(Fig 5B)**. To study the Mco10 protein levels we successfully generated a Mco10-specific antibody using the Mco10 fragment that differs from Atp19 (see Materials and Methods). Western blot analysis of total protein extracts showed that Mco10 protein levels do not change as compared to wild type cells when Atp19 or the dimer specific subunits Atp20 and Atp21, Atp14 or Atp18 were deleted (**Fig 5C, Appendix Fig S4)**.

**Figure 5.**
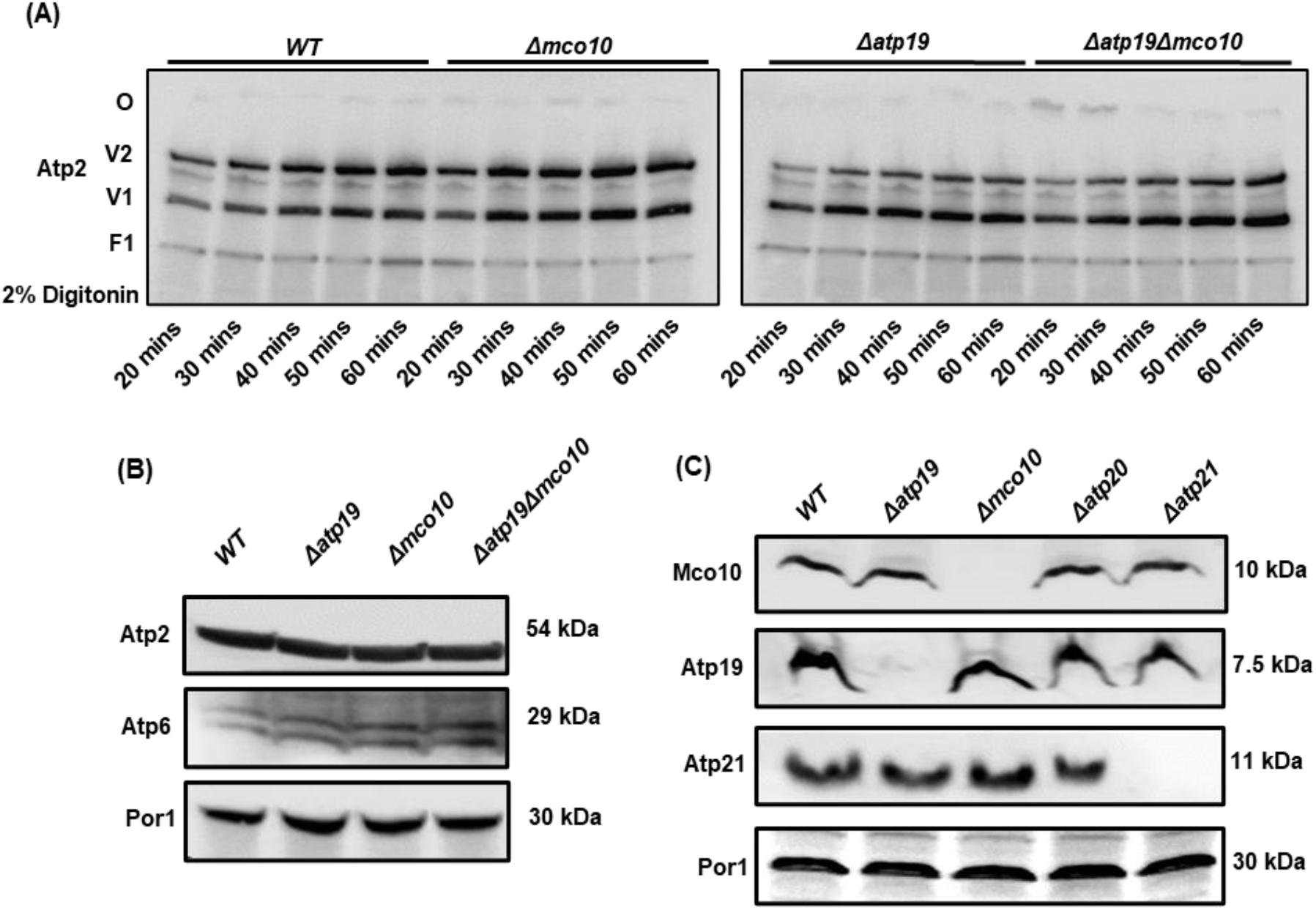
Mco10 deletion makes ATP synthase more susceptible to digitonin extraction. **A**. BN-PAGE analysis of ATP synthase complexes extracted from 500 pg mitochondria by incubation in extraction buffer containing 2 % digitonin for indicated time. The dimers (V2), monomers (V1) and free F1 domain are visualized by western blot using anti-Atp2 antibody. **B**. Western blot analysis of ATP synthase subunits in total protein extracts from *ΔmcolO, Δatp19* and *Δatp19Δmco10* mutants grown in YPGIyA medium. **C**. Steady-state levels of Mco10, Atp19 and Atp21 in indicated mutants determined with antibodies specific to these subunits. Pori protein level is used as loading control.

**Figure EF3.**
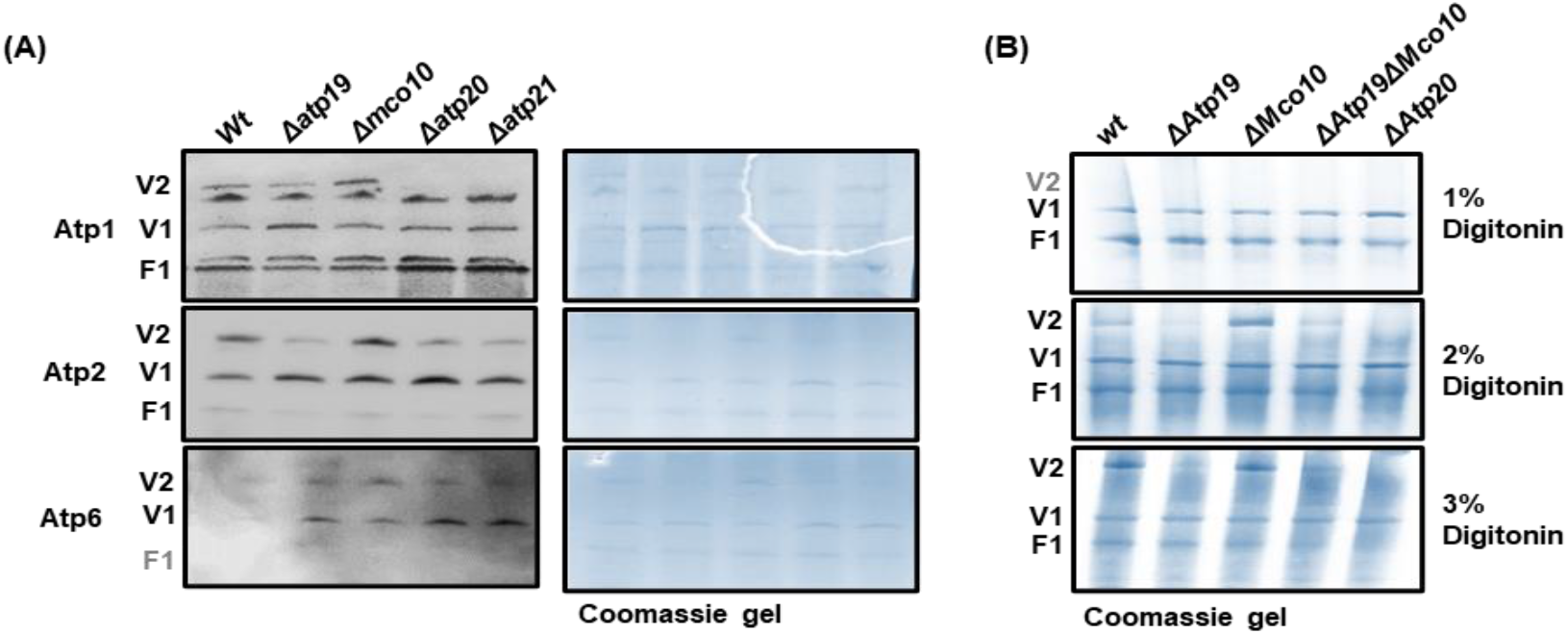
Mco10 deletion makes ATP synthase more susceptible to digitonin extraction. **A**. Monomers and dimers of ATP synthase from the wild type and indicated mutants were extracted by 2 % digitonin and western blot was performed using anti-Atp1, anti-Atp2 and anti-Atp6 antibody. **B**. Monomers and dimers extracted using 1 %, 2 % or 3 % digitonin and visualized by Coomassie staining.

### Mco10 mainly associates with the ATP synthase monomer

To confirm the Mco10 association with the ATP synthase we extracted its complexes, separated them in native gels and processed by Western blotting with anti-Mco10 antibody. Unfortunately, the anti-Mco10 antibody (as well as anti-Atp19 antibody used in this study) could not recognize the protein specifically in the ATP synthase monomer/dimer complexes - probably the peptide used for immunization is not exposed in the native Mco10 or Atp19 in the whole complex **(Appendix Fig S5A)**.

We then again applied the 2D-BN-SDS PAGE technique as done previously with mass spectrometric analysis of the ATP synthase monomers and dimers. Western blot with anti-Mco10 antibody revealed that Mco10 is indeed present with the complex, but mainly in the monomer similarly in wild type as well as in *Δatp19, Δatp20 and Δatp21* mutants **(Fig 6 and Appendix Fig S5B)**. Consistently, the Atp19 is present mainly in the dimer of ATP synthase. Because Mco10, Atp19, Atp21 and Atp20 were detected in both the monomers and dimers in mass spectrometric analysis we applied the emPAI calculations, which gives a rough estimation of the protein’s abundance. In accordance to emPAI, the Atp19 and Atp21 were respectively 16 and 100 times more abundant in the dimer when compared with the monomer while the Mco10 was 3 times more abundant in the monomer when compared with the dimer **(Appendix Fig S5C)**.

**Figure 6.**
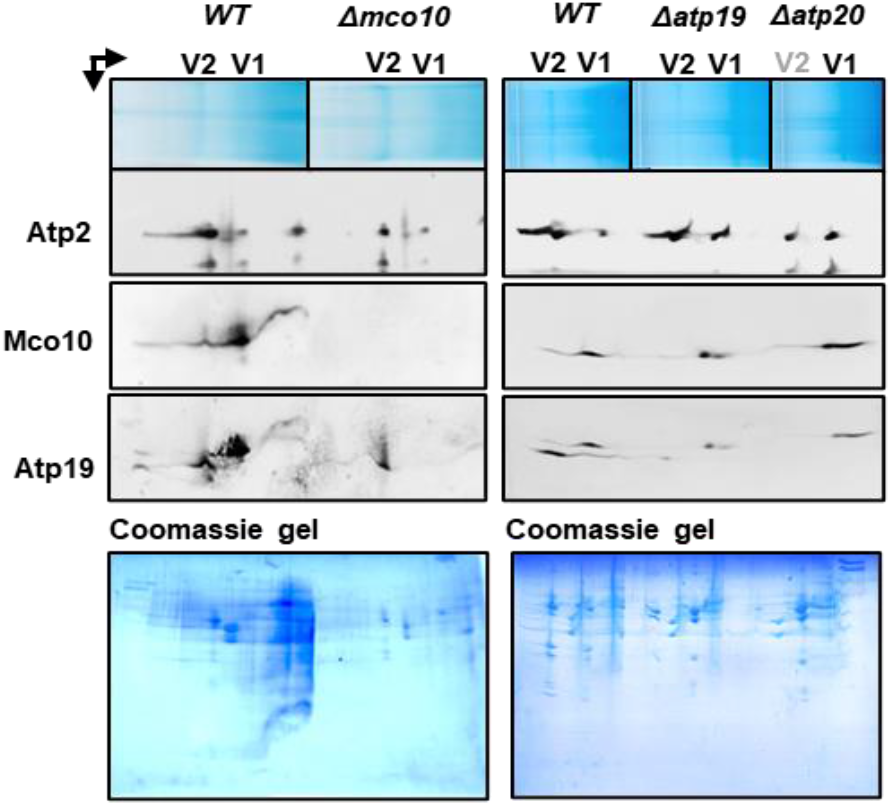
Mco10 is mainly present in the monomer of ATP synthase. The two-dimensional BN-SDS-PAGE gel separation of monomer and dimer subunits. After migration proteins were transferred to PVDF membrane and visualized by western blotting with respective antibodies. The upper panel shows the separation of the monomers and dimers in the first dimensional native gel. The lower panel shows the Coomassie stained second dimensional gels.

### Deletion of Mco10 affects calcium homeostasis and yPTP induction

Previous experiments showed that Mco10 protein is present in ATP synthase monomer and does not impact the ATP synthase activity and stability. We hypothesized then that Mco10 may be involved in the ATP synthase role in permeability transition. As transient PTP opening functions in calcium homeostasis we tested the survivability of wild type, *Δatp19, Δmco10, Δatp19Δmco10* and *Δatp21* (as a control, as lack of subunit *e* delays the pore opening in yeast (Carraro et al., 2014)) under high concentration of calcium in the media. Drop test analysis showed that at 1 M Ca^2+^ in the media, growth of *Δmco10, Δatp19Δmco10* and *Δatp21* were significantly more affected when compared to the wild type or *Δatp19* strain **(Fig 7A)**. This indicated that Mco10 deletion may affect yeast permeability transition pore (yPTP) and cell death.

**Figure 7.**
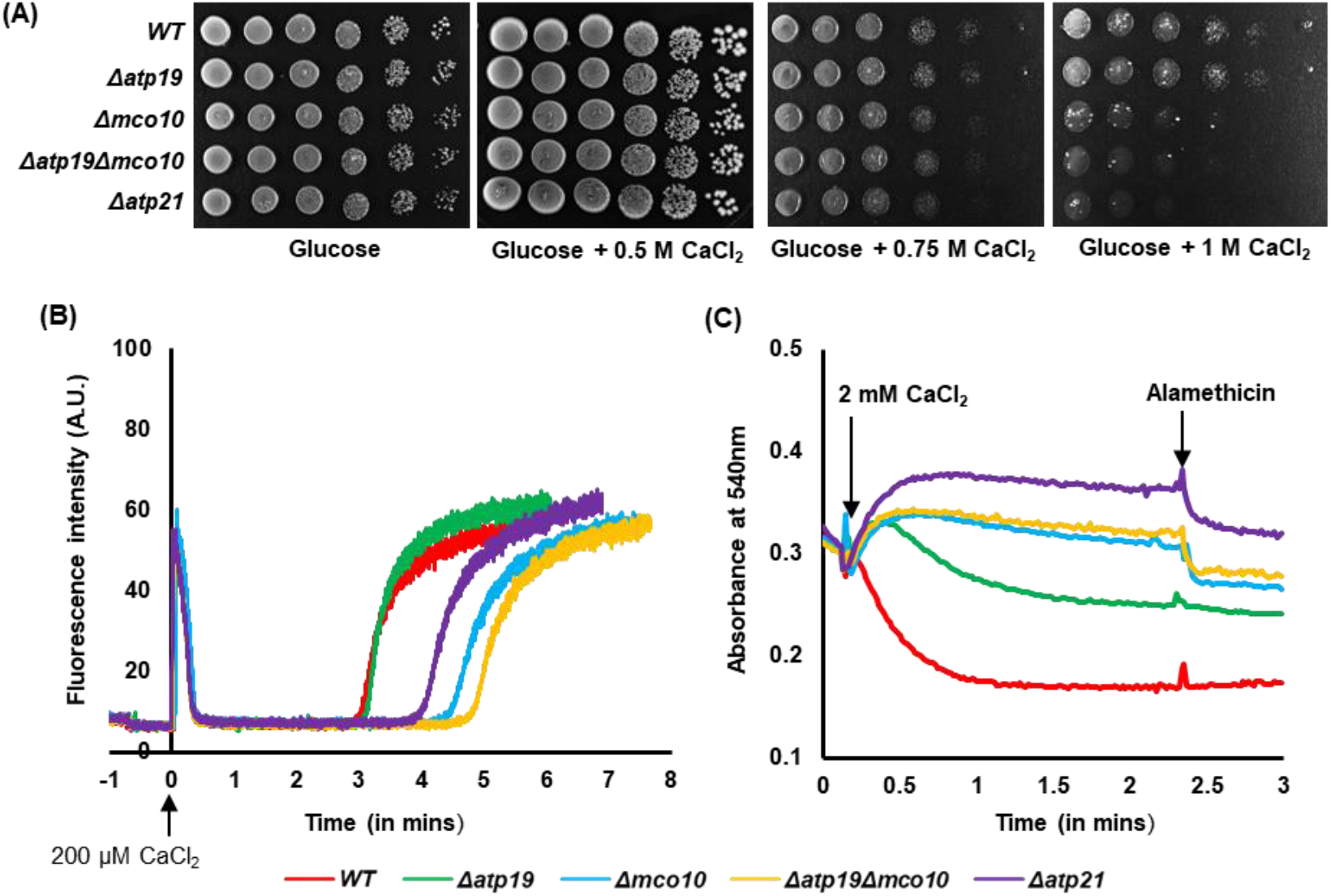
Mco10 modulates the yPTP. **A**. Cells from the indicated strains grown in glucose pre-cultures were serially diluted, spotted on rich glucose plates supplemented with 0.5 M, 0.75 M and 1 M CaCI_2_ and incubated at 28 “C. Plates were photographed after four days of incubation. **B**. The yPTP induction time measured in the CRC experiment. 1 mg of mitochondria was added to CRC buffer containing the calcium ionophore ETH129 and the calcium green-5N calcium indicator, and 200 pM CaCI_2_ was added at time 0. The experiment was monitored till the rapid increase of the calcium in the buffer. Traces are representative of at least 3 independent experiments **C**. Mitochondrial swelling assay. Swelling was induced by the addition of 2 mM CaCI_2_ to mitochondria suspended in swelling buffer containing the ETH129 and measured as the decrease in absorbance at 540 nm. Alamethicin (10 pM) was added after 2.5 minutes of swelling for normalization of the experiment. The representative plates and curves are shown.

To further confirm the role of Mco10 in yPTP, we analyzed the rate of yPTP opening after high calcium addition by measuring the time necessary for calcium release into the buffer using the calcium retention capacity (CRC) method. In this protocol, Ca^2+^ release occurs due to the depolarization of the membrane and therefore is compatible with the opening of a low conductance PTP (Niedzwiecka et al., 2020). Mitochondria were loaded with 200 µM CaCl_2_, which was completely taken up into the matrix via the calcium ionophore ETH129. After 3-5 minutes, calcium was released into the buffer through the yPTP in the mitochondria, which was measured by the increase in florescence of Calcium Green, a cell impermeable calcium ion concentration indicator. In the wild type and *Δatp19* mitochondria the release of calcium occurred after 3 minutes in these experimental conditions. As expected, the absence of Atp21 decreased Ca^2+^ sensitivity of the pore which leads to increase in the time required to open the pore when compared with the wild type mitochondria (Carraro et al., 2018). However, lack of Mco10 increased the yPTP opening time even longer than lack of Atp21, indicating that deletion of Mco10 significantly affects Ca^2+^ sensitivity of the yPTP. Deletion of *Δatp19* had no impact on the time of the pore opening. But in the *Δatp19Δmco10* double mutant, time of Ca^2+^ release was prolonged which further confirms our finding **(Fig 7B)**. The experiment was also performed with pulses of 10 µM CaCl_2_ added at an interval of 20 secs until the mitochondria stops the intake of Ca^2+^ ions and finally release it into the buffer. The release of calcium was similarly slower when Mco10 was deleted under these conditions **(Appendix Fig S6)**.

Next, we measured the swelling of mitochondria induced by high (2 mM) Ca^2+^ concentrations and caused by the diffusion of sucrose into the matrix. This is only possible when a large channel is formed. In this experiment, wild type mitochondria were able to form such a channel and swelled shortly after calcium addition. As expected, swelling was significantly blocked in *Δatp21* mutant mitochondria. Lack of Mco10 affected swelling similarly like lack of Atp21, while lack of Atp19 had moderate impact on mitochondrial swelling. Mitochondria from *Δatp19Δmco10* double mutant was also unable to swell properly **(Fig 7C)**. Thus, we conclude that Mco10, a new subunit of ATP synthase in *S. cerevisiae* is critical for calcium induced yPTP.

## Discussion

So far three supernumerary subunits of the mitochondrial ATP synthases were identified in different experimental approaches and their presence in the enzyme dimeric structures were confirmed (Arnold et al., 1998; Arselin et al., 2004; Wagner et al., 2010). These three subunits: *e, g* and *k*, fulfil the structural role in the stabilization of the enzyme dimers (Carraro et al., 2018; Habersetzer et al., 2013; Wagner et al., 2010). An additional fungi specific subunit *l* was co-purified with the ATP synthase of *S. cerevisiae* and *P. angusta* and their homologues were fund in fungi genomes but there is no evidence that the protein is a subunit of ATP synthase, nor any indication of its biological function (Liu et al., 2015). In our effort to purify *S. cerevisiae* ATP synthase and in-depth analysis of small proteins that interact with it, we identified Mco10 to be mainly associated with the monomer. This protein caught our attention due to its similarity to subunit *k* and the identification of phosphorylation at Ser-53 of Mco10 in mass spectrometry analysis. The phosphorylation of S56, S57 or S59 of Mco10 was also previously reported in a large-scale study of mitochondrial phosphoproteome (Renvoise et al., 2014). These post-translational modifications were found on residues of the central hydrophilic region of Mco10. The Mco10 peptide was not found in the crystal structures of the yeast ATP synthases in previous works with fungal ATP synthases, what questions whether Mco10 and its homolog in *P. angusta* is the subunit of ATP synthase (Guo et al., 2017; Hahn et al., 2016; Srivastava et al., 2018; Vinothkumar et al., 2016).

Lack of both proteins affected different functions of ATP synthase. In accordance to the published data, we found that Atp19 is needed for stabilization of the dimers of ATP synthase while Mco10 is not (Fig. 5A). The lack of Atp19 impacts the ATP synthesis activity of ATP synthase while lack of Mco10 has minor effect (Fig. 3C). The oligomycin sensitivity of ATP synthesis was however increased by lack of both Atp19 and Mco10 - although to different extend, arguing that Mco10 is indeed attached to the enzyme and modulates oligomycin effect on the enzyme activity **(Fig. 3B**,**F)**. Oligomycin binds to the *c*-ring subunits that are in contact with the proton half channel formed by subunit *a* (Nagley et al., 1986; Sebald et al., 1979; Symersky et al., 2012). We propose that Mco10 facilitates oligomycin binding or blocking of the *c*- ring rotation (see below). In *Δatp19*, more of Mco10 may be present on ATP synthase complexes therefore these complexes are more efficiently inhibited by this drug.

From the crystal structures of *S. cerevisiae* and bovine ATP synthases, Atp19 attaches with the complex with its N-terminal helix in a V-shaped groove formed by helix 4 and 5 of subunit Atp6/*a*. Basing on the conservation of the N-terminal fragments of Mco10 and Atp19 sequence and structures in *S. cerevisiae* and in other yeast, we propose that Mco10 and Atp19 might be present in the same position in the ATP synthase F_O_ domain (Guo et al., 2017). To support this hypothesis, we reconstructed the Fo domain based on the crystal structure (PDB: 6B8H) with AlphaFold2 predicted structures of each of its subunits and replaced the predicted AlphaFold2 structure of Atp19 with that of Mco10 in this model. Surprisingly, Mco10 accommodates very well in the crystal structure with least steric hindrance with the neighboring subunit Atp18/*i*, Atp6/*a* or the *c*-ring **(Fig EF4)**. The fact that this binding pocket is in close proximity to the proton half channel and oligomycin binding site in the *c*-ring supports our hypothesis that oligomycin blocks more efficiently the rotation of the *c*-ring primed by Mco10. Although Mco10 was identified in the both monomers and dimers by mass spectrometry analysis it is more abundant in the monomer of ATP synthase in western blot analysis. It was previously shown that Atp19, Atp20 and Atp21 subunits ‘prime’ the monomers before the dimers are formed (Wagner et al., 2010). Basing on the increased sensitivity to oligomycin of the ATP synthase in *Δatp19* cells and the proposition of Symersky et al., 2012 that oligomycin may preferentially bind to the *c*-ring of the ATP synthases that are damaged and uncoupled (based on the previous findings that oligomycin increases phosphorylation capacity of EDTA or alkali-treated non-phosphorylating sub-mitochondrial particles (Fessenden & Racker, 1966; Lee & Ernster, 1965)), we further propose that the ATP synthase where Mco10 binds may be uncoupled/damaged physiologically.

**Figure EF4.**
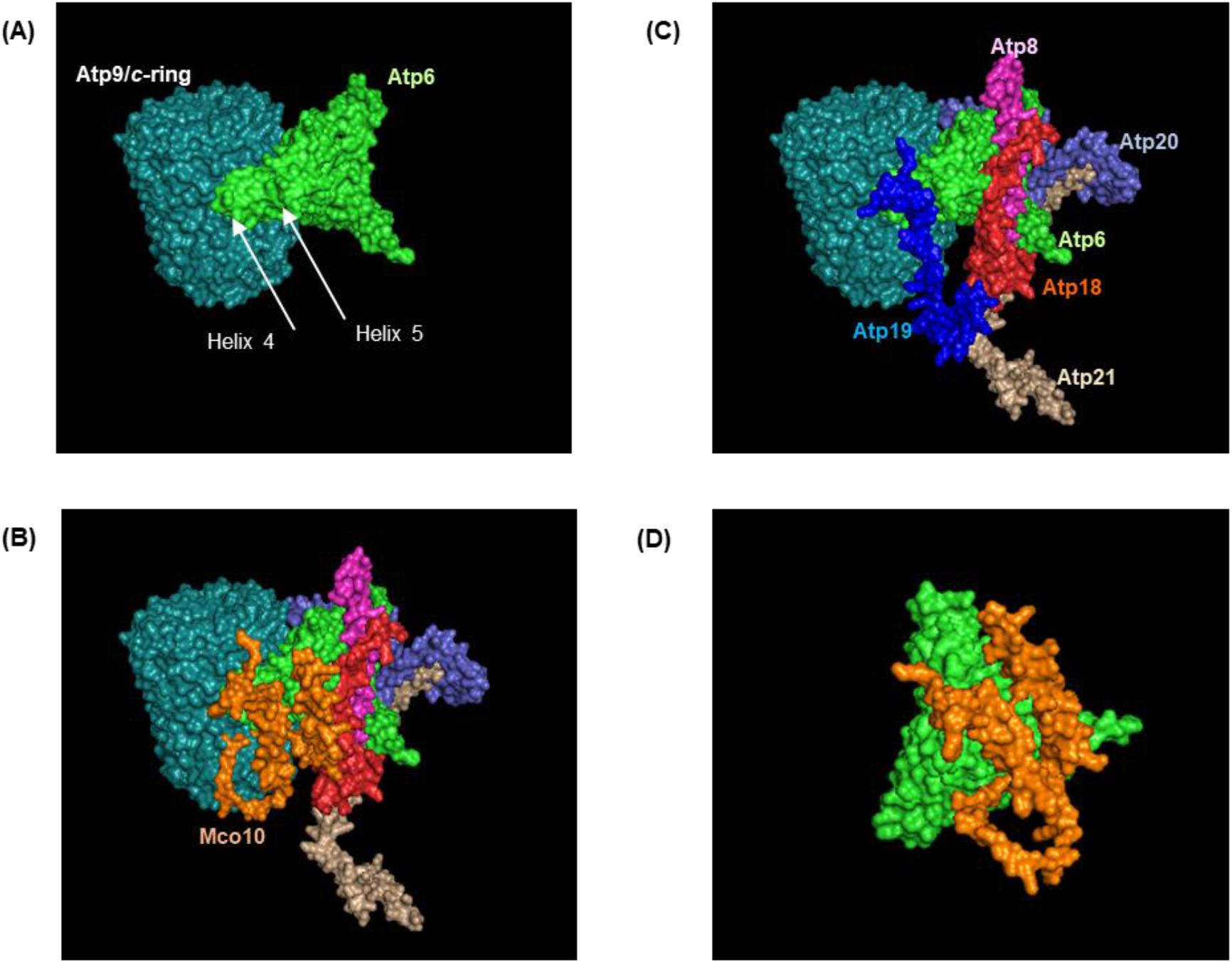
Remodeling of ATP synthase Fo region with AlphaFold2 predicted structures of the Fo subunits using PDB 6B8H as template and visualized in PyMOL. **A**. Orientation of Atp6 with respect to the Atp9/c-ring. The groove formed by helix 4 and 5 of Atp6 where Atp19 binds are indicated. **B**. Partial reconstruction of the Fo region showing Atp19 (deep blue), Atp18 (red), Atp6 (green), Atp21 (golden), Atp20 (light violet). **C**. Atp19 is replaced by Mco10 (orange) in the model. **D**. A close-up view of the orientation of the two helices ofMcolO with respectto its alignment with Atp6.

Another very important role of Mco10, but not Atp19, for mitochondrial physiology is yPTP regulation by calcium and possibly regulation of calcium homeostasis. It is documented, also in yeast, that yPTP is delayed in the absence of the dimerization subunits *e* and *g* indicating that ATP synthase dimers are needed for channel formation (Carraro et al., 2014; Chinopoulos, 2018; Giorgio et al., 2013). Our data show that among the supernumerary ATP synthase subunits, only subunit *k* does not appear to be involved in formation of the megachannel. Importantly, cells lacking Mco10 but not Atp19, similarly like those lacking Atp21/*e* that is necessary for yPTP induction, grow slower in high concentration of calcium in the medium. Calcium homeostasis has a direct link with reactive oxygen species (ROS) generation and increase in calcium cytosolic concentration leads to cell death through mitochondrial permeability transition (Bertero & Maack, 2018; Gorlach et al., 2015). Thus, the decreased viability of Mco10 or Atp21 lacking cells in media supplemented with calcium indicates that indeed yPTP contributes to calcium buffering by its pumping into the mitochondria, as we suggested previously (Niedzwiecka et al., 2018). The yeast mitochondria do not possess a known calcium transporter to the mitochondrial matrix (Bradshaw & Pfeiffer, 2006; Carafoli et al., 1970; Carraro & Bernardi, 2016). The Sec61 translocation complex, subunits of which were identified in the interactome analysis of both yeast and human ATP synthase, can also contribute to this process, as it was shown previously to form ubiquitous Ca^2+^ leak channels in the endoplasmic reticulum (Lang et al., 2011).

How ATP synthase forms the channel is not understood yet. One hypothesis proposes that dimerization of ATP synthase is essential for generation of the pore and pore is formed upon dissociation of the monomers (Chinopoulos, 2017; Giorgio et al., 2017) or alternatively by the F_O_ part of the dimer (Bernardi et al., 2021). Another model proposes that ATP synthase monomer constitutes the pore forming part and its structural rearrangements within the F_O_ sector that perturb subunit *e* and may exert a pulling action dragging lipids out of the *c*-ring permitting transduction through the *c*-ring (Alavian et al., 2014; Bernardi et al., 2021; Bonora et al., 2013). The ability of monomers to form PTP was also shown in mammalian liver and heart mitochondria (Bonora et al., 2017; Mnatsakanyan et al., 2019). We propose that in yeast upon the PTP induction, the subpopulation of ATP synthase ‘primed’ by Mco10 forms the PTP. The hydrophilic middle region of Mco10 may interact with the *c*-ring that help in maintaining a ‘open’ channel conformation (Mnatsakanyan et al., 2022). When Mco10 is deleted, the *c*-ring cannot maintain an open channel that leads to inhibition of PTP. The fact that Atp19 does not directly interacts with the *c*-ring and its deletion alone does not inhibit yPTP but it is inhibited in *Δatp19Δmco10* double mutant, further proves the association of Mco10 with ATP synthase complex involved in yPTP.

The important question that still remains unanswered is whether monomer of human ATP synthase has a functional homolog of Mco10. Although our small molecule interactome analysis did identify potential unknown modulators of yeast ATP synthase in which future research can be directed, we failed to identify such an unknown subunit/s in the human ATP synthase interactome. Based on important Mco10 role for ATP synthase function and conservation of ATP synthase structure and function in yeast and human, it is likely that similar mechanism of PTP modulation by a specific protein that only associates with a subpopulation of ATP synthase complexes does exists in mammals. This can well be an isoform of a known ATP synthase subunit. If a functional homolog of Mco10 exists in human ATP synthase, and fulfils similar function like in yeast in modulation of PTP but not the ATP synthesis activity or dimerization of the complex, it would be an ideal target for the drugs screen for treatment of neurodegenerative diseases (Garone et al., 2022; Mnatsakanyan et al., 2022; Ortiz-Gonzalez, 2021).

## Material and methods

### Yeast strains and growth media

Strains of *Saccharomyces cerevisiae* used in the study were BY4741 (*MATa his3Δ1; leu2Δ0; met15Δ0; ura3Δ0*) and isogenic *Δatp19, Δatp20, Δatp21* and *Δmco10* (Euroscarf collection). The double mutant *Δatp19 Δmco10* was constructed by crossing the single mutants in the opposite mating sign and tetrad dissection. MR6 (*MATa ade2-1 his3-1,15 leu2-3,112 trp1-1 ura3-1 arg8::HIS3*) expressing Atp6- HA-6His was a gift from prof. Alexander Tzagoloff (Columbia University, NY, USA). Strains were grown in rich YPGA medium (1% Bacto yeast extract, 1% Bacto peptone, 2% glucose, 40 mg/l adenine), YPGlyA medium (1 % Bacto yeast extract, 1 % Bacto peptone, 2 % glycerol, 40 mg/l adenine) or W_0_ complete minimal medium (6.7 % yeast nitrogen base w/o amino acids and 2 % glucose, supplemented with appropriate drop-out amino acid stock (Sunrise)) at 28°C or 36°C with shaking at 200 rpm. The liquid media were solidified by addition of 2% Bacto agar (Difco, Becton Dickinson). For mitochondria isolation, strains were grown in rich YPGlyA medium to an OD_600_ of 4. An OD_600_ of 1 corresponds to 1.2 × 10^7^ cells/ml. G418 sulphate were added to media at a concentration of 200 µg/ml when required.

### Purification of ATP synthase complexes from mitochondria

The strain expressing Atp6 subunit C-terminally tagged by HA-6xHis in the mitochondrial genome was characterized previously to cause no damage to the ATP synthase structure and activities (Rak et al., 2009). This strain was used to pull down the whole ATP synthase complex by Ni-NTA agarose beads. Briefly, 5 mg of mitochondria were centrifuged and suspended in 1 mL of sonication buffer (250 mM sucrose, 50 mM NaH_2_PO_4_, 5 mM 6-aminocaproic acid, 1 mM EDTA, pH 7.5, protease inhibitors cocktail tablet (Roche), 1 µM PMSF) and sonicated 6 times 10 sec, with 10 sec intervals on ice. After centrifugation at 6000 x g 10 minutes at 4°C, supernatant was ultracentrifuged at 268526 x g for 1 hour (Thermo Scientific™ Sorvall™ WX ultracentrifuge, TFT80.2 rotor). The pellet was washed twice with the sonication buffer without EDTA (without suspending it) and then suspended with the use of the potter in 500 µl MP extraction buffer (150 mM potassium acetate, 10 % glycerol, 2 mM 6-aminocaproic acid, 30 mM HEPES, pH 7.4, 1 % N-dodecyl-β-maltoside, 2 mM PMSF and protease inhibitors cocktail tablet, EDTA-free, Roche). After 20 minutes incubation on ice, the membranes were centrifuged for 30 mins at 21950 x g, 4 °C and the extract was incubated with 200 µL of the Ni-NTA agarose washed previously by Binding buffer (50 mM NaCl, 10 % glycerol, 10 mM imidazole, 20 mM NaH_2_PO_4_, pH=7.9, 0,1% n-dodecyl-β-maltoside, 2 mM PMSF, protease inhibitors cocktail tablet) for overnight. Next day the beads were washed twice with the Binding buffer, then suspended in 400 µL of Binding buffer, dosed and after addition of 100 µL of 5x Laemmli sample buffer, boiled 5 minutes. The 50 µg of the extract and 2 µg of bead eluate were loaded on the 15 % SDS-PAGE gel. Then the gel was stained with Coomassie blue or silver staining according to manufacturer’s protocol (Pierce sliver stain kit, Thermo Fisher Scientific) to visualize the proteins.

### Isolation of mitochondria from HEK293T cells

HEK293T (ATCC) cells were maintained in media composed of DMEM (high glucose; Biowest), sodium pyruvate (Thermo Fisher Scientific), stable glutamine (Biowest), non-essential amino acids (Thermo Fisher Scientific), 10% fetal bovine serum (EurX), and penicillin/streptomycin solution (Thermo Fisher Scientific). Cells were cultured to confluence on six 9cm plates. Mitochondria were isolated by differential fractionation previously described (Spelbrink et al., 2000) with minor adjustments. Briefly, cells were washed 2x with ice-cold PBS. The PBS buffer was removed and the cells were scraped in ice-cold NKM buffer (1 mM Tris-HCl pH 7.4, 130 mM NaCl, 5 mM KCl, 7.5 mM MgCl2) and incubated on ice for 5 minutes. Then 100 μl of 10xHomB buffer (225 mM mannitol, 75 mM sucrose, 10 mM HEPES-NaOH pH 7.8, 10 mM EDTA) per 1 ml of homogenate with protease and phosphatase inhibitors was added and homogenized using a Dounce homogenizer with nuclei release and integrity monitored microscopically. The nuclear fraction was separated using two rounds of centrifugation at 900xg for 5 mins at 4°C. The supernatant was then centrifuged at 10000xg for 10 mins to isolate crude mitochondrial pellet. The pellet was suspended in Blue Native protein extraction buffer, dosed and ATP synthase complex from 400 μg of protein was extracted with 2 % Digitonin. The dimers and monomers were separated in 3-12 % gradient gel followed by SDS-PAGE separation in second dimension and bands from 5-20 kDa region from Coomassie stained gel from the monomer and dimer were subjected to mass spectrometric analysis.

### Production of Mco10 antibody

The Mco10 specific antibody was generated using a peptide KDDDVVKSIEGFLNDLEKDTRQDT produced synthetically identical to the Mco10 fragment from 64 to 83 amino acids residues by immunizing rabbit and isolating the affinity purified polyclonal antibody from Davids Biotechnologie GmbH (Germany).

### Two Dimensional BN-PAGE/SDS-PAGE and Western blotting

Two-dimensional gel electrophoresis was based on the protocol of Schamel (Schamel, 2008) with slight modifications. Briefly, the ATP synthase complexes were liberated from inner mitochondrial membrane of isolated mitochondria by incubation with 1-2 % digitonin in extraction buffer (30 mM HEPES, 150 mM potassium acetate, 12 % glycerol, 2 mM 6-aminocaproic acid, 1 mM EGTA, protease inhibitor cocktail tablets EDTA-free (Roche), pH 7.4) for different time intervals up to 60 mins and separated using NativePAGE™ 3-12 % Bis-Tris Gels (Thermo Fisher Scientific) to separate monomeric and dimeric ATP synthase complexes (Schagger & von Jagow, 1991). For second dimensional analysis the lanes were cut from the gel and placed in SDS-PAGE running buffer (25 mM Tris, 192 mM Glycine, 0.1 % SDS, pH 8.3 with 1 % β-mercaptoethanol), heated in a microwave for 10 secs and incubated for another 10 mins in a shaker. The gel strips were then loaded on the top of a 16 % SDS-PAGE gel, and electrophoresis was conducted under denaturing conditions. Then the gel was stained with Coomassie blue or silver staining and bands cut-off were analyzed by mass spectrometry or proteins from the gel were then transferred into PVDF or nitrocelluse membranes using iBlot system (Thermo Fisher Scientific). For SDS-PAGE analysis of steady state level of proteins, yeast cells were disrupted by alkaline lysis with NaOH/TCA (Kushnirov, 2000). Western blot analysis was performed using the polyclonal rabbit anti-Mco10 antibody or anti-ATP synthase subunit antibodies (gifts from Marie-France Giraud, Bordeaux, France and Martin van der Laan, Germany).

### Mass spectrometry analysis

The proteins in the 5-20 kDa region from SDS-PAGE gels were cut, de-stained and subjected to in-gel tryptic digestion. The tryptic digested peptides were then independently analyzed by LC/MS system at the IBB PAS Mass Spectrometry Facility unit using Evosep One (Evosep Biosystems, Odense, Denmark) coupled to a Orbitrap Exploris 480 mass spectrometer (Thermo Fisher Scientific, Bremen, Germany) via Flex nanoESI ion source (Bache et al., 2018) or by Thermo EASY-nLC 1000 system interfaced to ThermoFisher LTQ Orbitrap Elite and Velos. Peptides were then loaded on a trapping column and eluted over a C18 75μm (Elite/Velos) or 150 µm (Exploris 480) analytical column at 250 nL/min. The mass spectrometer was operated in data-dependent mode. MS1 scan range of 300 to 1600 m/z was selected and top 40 precursors within an isolation window of 1.6 m/z were considered for LC/MS analysis. Data was acquired in positive mode with a data-dependent method using the following parameters: the raw data were then searched using an inhouse copy of Mascot with the following parameters: Enzyme: Trypsin, Database: SGD_new (6,713 sequences; 3,019,540 residues) or Sprot_02 (567,483 sequences; 204,940,973 residues) with Carbamidomethyl (C) as fixed modification and Oxidation (M), Acetyl (N-term, K), Phospho (S, T, Y), GlyGly (K) as variable modifications, Mass values: Monoisotopic, Peptide Mass Tolerance: ±5-10 ppm, Fragment Mass Tolerance: ± 0.01-0.1 Da, Max Missed Cleavages: 2 and rejected all proteins found in cRAPome database during analysis.

### Measurement of oxygen consumption, ATP synthesis and membrane potential

Mitochondria were isolated from cells grown in YPGlyA medium at 28°C by enzymatic method according to the protocol described previously (Guerin et al., 1979). For all assays, they were diluted to 75 µg/ml in respiration buffer (10 mM Tris-maleate pH 6.8, 0.65 M mannitol, 0.35 mM EGTA, and 5 mM Tris- phosphate). Oxygen consumption rates were measured using a Clarke electrode adding consecutively 4 mM NADH (state 4 respiration), 150 µM ADP (state 3) or 4 µM carbonyl cyanide m-chlorophenylhydrazone (CCCP) (uncoupled respiration), as described previously (Rigoulet & Guerin, 1979). Complex IV activity was measured by first adding to mitochondria and CCCP to uncouple respiration. Then, simultaneously 100 µM ascorbate and 3 µM N,N,N′,N′-Tetramethyl-p-phenylenediamine (TMPD) were added and oxygen consumption rates measured. The rates of ATP synthesis were determined under the same experimental conditions with 750 µM ADP; Every 15 seconds, 100 µl aliquots were withdrawn from the oxygraph cuvette and added to 50 µl of the 3.5 % (w/v) perchloric acid and 12.5 mM EDTA solution already prepared in the tubes to stop the reaction. The samples were then neutralized to pH 6.5 by the addition of KOH and 0.3 M MOPS. The synthetized ATP was quantified using a luciferin/luciferase assay (Kinase-Glo Max Luminescence Kinase Assay, Promega) in a Beckman Coulter Paradigm plate reader. The participation of F_1_F_O_-ATP synthase in ATP production was assessed by measuring the sensitivity of ATP synthesis to oligomycin (3 μg/ml). The specific ATPase activity at pH 8.4 of non-osmotically protected mitochondria was measured using the procedure previously described (Somlo, 1968). The oxygen consumption was quantified in nmol O_2·_min^-1^·mg^-1^, the ATP synthesis in nmol of ATP.min^-1^.mg^-1^ and ATPase activities in µmol Pi.min^-1^.mg^-1^. Variations in transmembrane potential (ΔΨ) were evaluated by monitoring the fluorescence quenching of Rhodamine 123 (0.5 μg/mL; λ_exc_ of 485 nm and λ_em_ of 533 nm) from mitochondrial samples (0.150 mg/mL) in the respiration buffer under constant stirring at 28°C using a Cary Eclipse Fluorescence Spectrophotometer (Agilent Technologies, Santa Clara, CA, USA) as described previously (Emaus et al., 1986).

### Measurement of the mitochondrial calcium retention capacity and swelling

A previously developed method for measurement of mitochondrial calcium retention capacity (described in detail in (Carraro et al., 2014)) was used to measure the time of yPTP opening after Ca^2+^ addition. Briefly, isolated mitochondria were diluted in CRC buffer (250 mM sucrose, 10 mM Tris-MOPS, 10 µM EGTA-Tris, 5 mM P_i_-Tris, 1 µM Calcium Green-5N (Thermo Fisher), 0.5 mg/ml BSA, pH 7.4) to a concentration of 1 mg/ml. The reaction was started by adding 1 mM NADH and 5 µM of Calcium ionophore ETH129. After equilibration for 1 min, 200 µM CaCl_2_ was added. The rapid increase in the fluorescence of Calcium Green-5N after sometime was attributed to the release of calcium ions from the mitochondrial matrix into the buffer, likely due to the opening of the permeability transition pore. Matrix swelling was evaluated by measuring optical density changes at 540 nm with a Parkin Elmer Lambda 925 UV-Vis spectrophotometer. Mitochondria (500 µg/ml) were suspended in 2 ml of swelling buffer (150 mM sucrose, 10 mM Tris-HCl, 2 mM KH_2_PO_4_, pH 7.4) and then, 2 mM CaCl_2_ and 10 µM alamethicin were added (Carraro & Bernardi, 2020).

### Phylogenetic tree construction

The first 25 amino acids of Mco10 or Atp19 of *S. cerevisiae* were used as templates to identify homologs of these proteins from the published fungal genomes using BlastP. The phylogenetic tree was constructed using representative example of homologs from the evolutionary branches of the phylum *Ascomycota* and *Basidiomycota*. The evolutionary history was inferred by using the Maximum Likelihood method and Poisson correction model (Felsenstein, 1981). The tree with the highest log likelihood (- 3887.59) is shown. Initial tree(s) for the heuristic search were obtained automatically by applying Neighbor-Join and BioNJ algorithms to a matrix of pairwise distances estimated using the Poisson model, and then selecting the topology with superior log likelihood value. This analysis involved 23 amino acid sequences. There was a total of 100 positions in the final dataset. Evolutionary analyses were conducted in MEGA X software (Kumar et al., 2018).

### Structural analysis

Multiple sequence alignment of Atp19 and Mco10 was performed using ClustalOmega (Madeira et al., 2022). Hydropathy plots were generated by ProtScale on the ExPASy Server using Kate and Doolittle for predicted proteins of *S. cerevisiae*. Homology modelling of the ATP synthase complex was based on the atomic model built in the cryo-electron microscopy density map of the *S. cerevisiae* ATP synthase, PDB: 6B8H (Guo et al., 2017), and Alphafold2 predicted structures of the subunits in *S. cerevisiae* (Jumper et al., 2021; Varadi et al., 2022). Structures of the homologs of Mco10 and Atp19 in *Candida albicans* and *Pichia angusta* were analyzed using the available structures in Alphafold2 database and also verified using ColabFold which offers an accelerated protein structure predictions by combining MMseqs2 with AlphaFold2 or RoseTTAFold (Mirdita et al., 2022). Structural visualization was carried out using PyMOL software.

### Data and Statistical analyses

To determine a high confidence small protein ATP synthase interactome, we emphasized more on manual validation due to poor performance of conventional algorithms which often rejects peptide fragments identified from these small proteins as false positive due to low score. Often, these are identified with a true single peptide hit but rejecting them makes the data underrepresented. To overcome this, we enriched the data using the following rules: all proteins identified with mol. wt ≥ 20 kDa were excluded from the analysis; proteins identified with single peptides were retained if identified in more than two independent analyses with protein coverage ≥ 10%. Most single peptide and low coverage identified proteins were further manually inspected to minimize false discoveries. Protein interaction network was constructed using STRING database with medium confidence level (Szklarczyk et al., 2019) and visualized using the STRING 106 plugin in Cytoscape. The identified proteins were classified manually or by Gene Ontology terms. Unless otherwise stated in the figure legends, each experiment was repeated at least three times. Data are presented as a representative experiment or as the average ± s.d. To assess significant differences with the control and test samples, Student’s t-test was used. All figures in this study were assembled using Microsoft Office PowerPoint or Adobe Photoshop (Adobe Inc.).

## Supporting information

Small Protein Interactome of S. cerevisiae ATP Synthase

Small Protein Interactome of Human ATP Synthase

Protein sequences of Atp19 and Mco10 homologs in different fungal genomes

Mascot Search result of proteins identified by LC/MS analysis

## Data Availability

The list of proteins identified in each mass spectrometry analysis from Database search using Mascot is available in **Dataset EV1**.

## Acknowledgements

We thank Prof. A. Tzagoloff for the MR6 ATP6-HisHA strain, Dr. Marie-France Giraud for the anti-ATP synthase subunits antibodies, Prof. Martin van deer Laan for anti-subunit *k* antibody, Prof. Adrianna Skoneczna for providing deletion strains from Euroscarf Library and Dr. Sampurna Datta for helping with cell culture. We also thank the IBB Mass spectrometry lab lead by Prof. Michal Dadlez and team for the help with mass spec analysis. This work was supported by a grant from the National Science Center of Poland (2018/31/B/NZ3/01117) to RK. The permission number for work with genetically modified microorganisms (GMM I) for RK is 01.2-28/201.

## Author Contributions

**CP:** Conceptualization; Data curation; Formal Analysis; Investigation; Methodology; Validation; Visualization; Writing – original draft; Writing – review & editing. **AW:** Data curation; Investigation; Writing – review & editing. **KN:** Investigation. **EB:** Investigation. **RK:** Conceptualization; Data curation; Formal Analysis; Funding acquisition; Investigation; Project administration; Resources; Supervision; Validation; Writing – original draft; Writing – review & editing.

In addition to the <u>CRediT</u> author contributions listed above, the contributions in detail are:

CP and RK conceived the project, CP designed all the experiments, performed the work, analyzed and interpreted the data, wrote the manuscript; AW performed membrane potential measurements, CRC and swelling experiments on native mitochondria and helped CP in conducting experiments, KN and EB performed the pull-down of ATP synthase experiments, RK supervised the whole work, analyzed and interpreted the data, wrote the final manuscript. All authors commented on the manuscript.

## Table and Dataset Legends

**Table EV1. Small ≤ 20 kDa proteins interactome of ATP synthase in *S. cerevisiae***. The proteins were identified from two different approaches using pulldown of ATP synthase by Atp6 tagged with HA-6His, or the monomers and dimers extracted by digitonin and separated in BN-PAGE or a further 2^nd^ dimensional SDS-PAGE identified by mass spectrometric analysis.

**Table EV2. Small ≤ 20 kDa proteins interactome of ATP synthase in HEK293T cell line. The** monomers and dimers were extracted from isolated mitochondria from HEK293T cells by digitonin and separated in a 2^nd^ dimensional BN-SDS-PAGE. Gel pieces from molecular weight ≤ 20 kDa were cut-off and proteins were identified by mass spectrometry.

**Table EV3. Protein sequences of** Atp19 and Mco10 homologs in different fungal genomes classified according to subphylum.

**Dataset EV1**. Mascot search result of proteins identified in each band by LC/MS analysis from all experiments performed in this study. The details of bands excised from gels are present in Appendix Fig S1.

