## Supplementary Appendix File for "ATP synthase interactome analysis identifies Mco10 – a new modulator of permeability transition pore in yeast"

**Small Protein Interactome analysis of ATP synthase identifies the uncharacterized ‘subunit’ Mco10 - a new modulator of permeability transition pore in *S. cerevisiae***

Chiranjit Panja*, Aneta Wiesyk, Katarzyna Niedźwiecka, Emilia Baranowska, Roza Kucharczyk*

Institute of Biochemistry and Biophysics, Polish Academy of Sciences, Warsaw, Poland

**Table of Content	Page no**

Figure S1 and S2	1

Figure S3 and S4	2

Figure S5	3

Figure S6	4

**(B)**

**(A)**

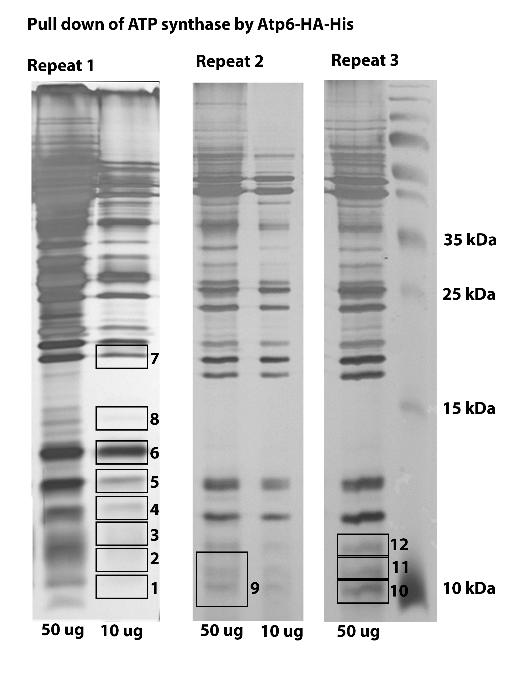

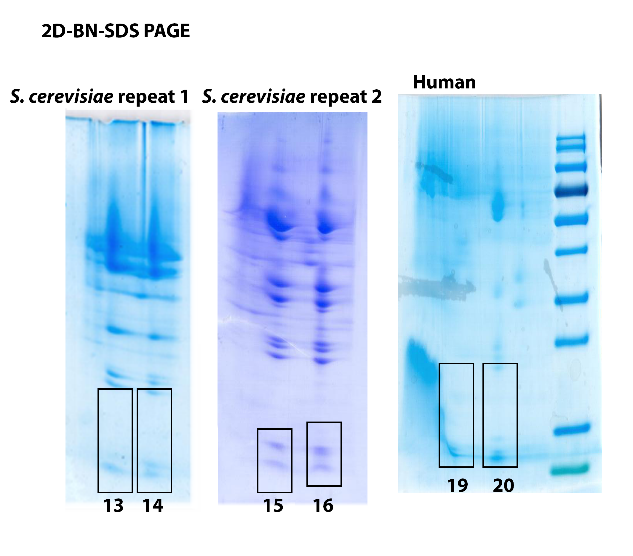

**Fig S1. A.** Repeat of pulldown experiments and excised gel regions used for mass spectrometric identification. The numbers correspond to the band numbers as described in Dataset EV1. **B.** Gel regions used for identification from 2D- BN-SDS-PAGE separation of monomers and dimers. Bands 17 and 18 (repeat 3) were cut from whole monomer and dimer from BN PAGE and not shown.

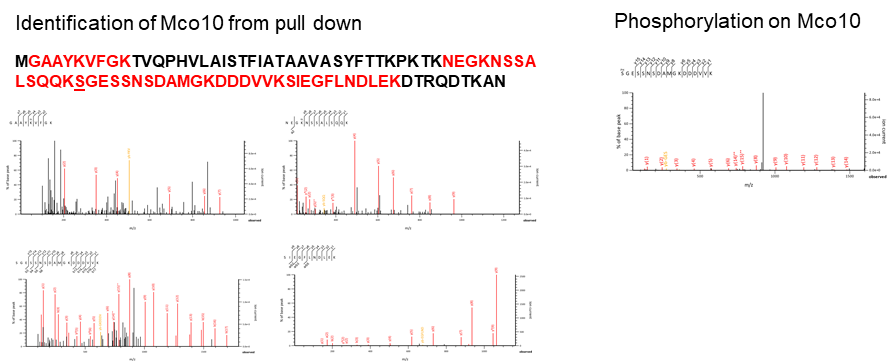

**(B)**

**(A)**

**Fig S2.** Identification of Mco10 in the ATP synthase interactome. **A.** The LC/MS spectra of four independent Mco10 peptides. Sequence marked in red were identified in the analysis. **B.** Phosphopeptide (phosphorylation at Ser53) of Mco10 identified in the LC/MS analysis.

**1**

**Fig S3.** Sequence alignment of Atp19 and Mco10 homologs in fungi. (1) and (2) represents Mco10 or Atp19 related homolog where two of them were present in the fungal genome.

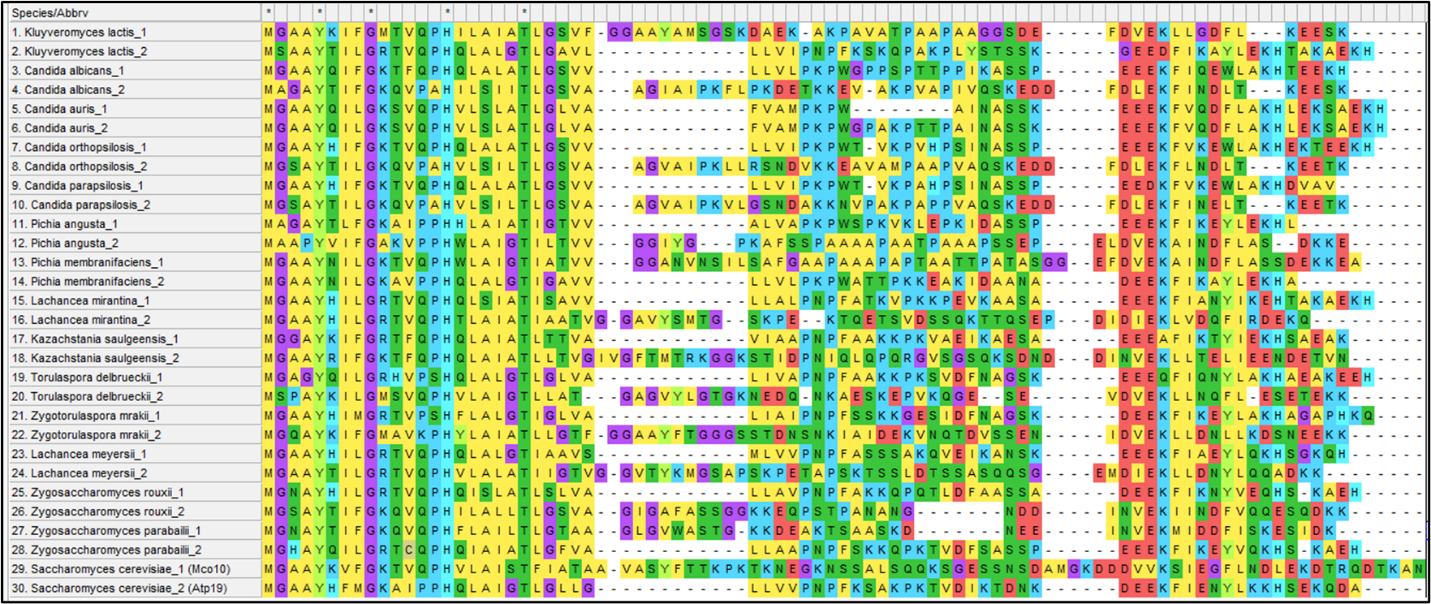

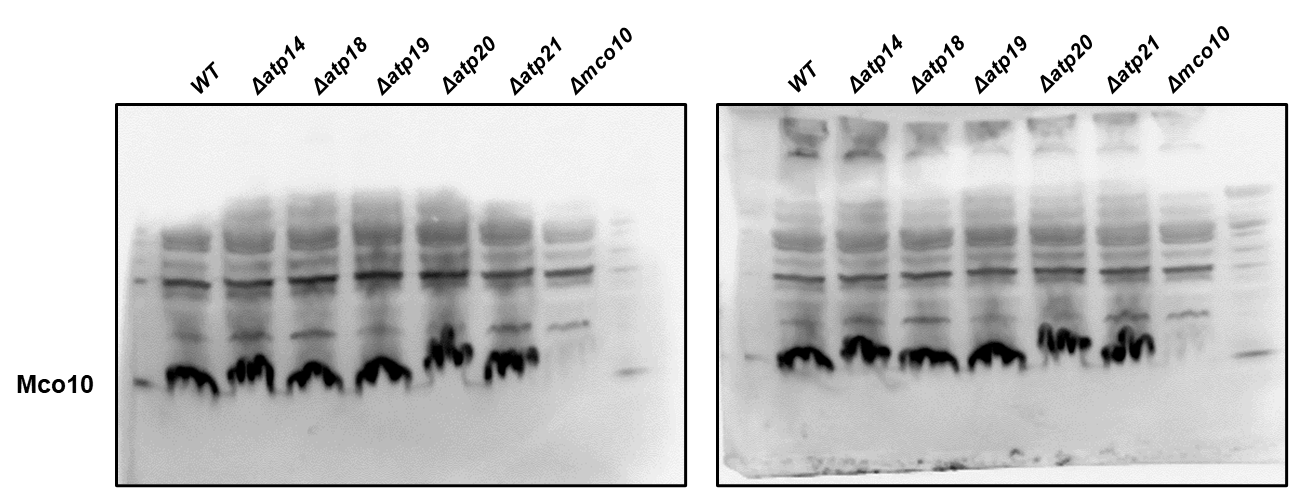

**Fig S4.** Steady-state levels of Mco10 in wildtype and indicated mutants determined with anti-Mco10 antibody. Two duplicate experiments are shown.

**2**

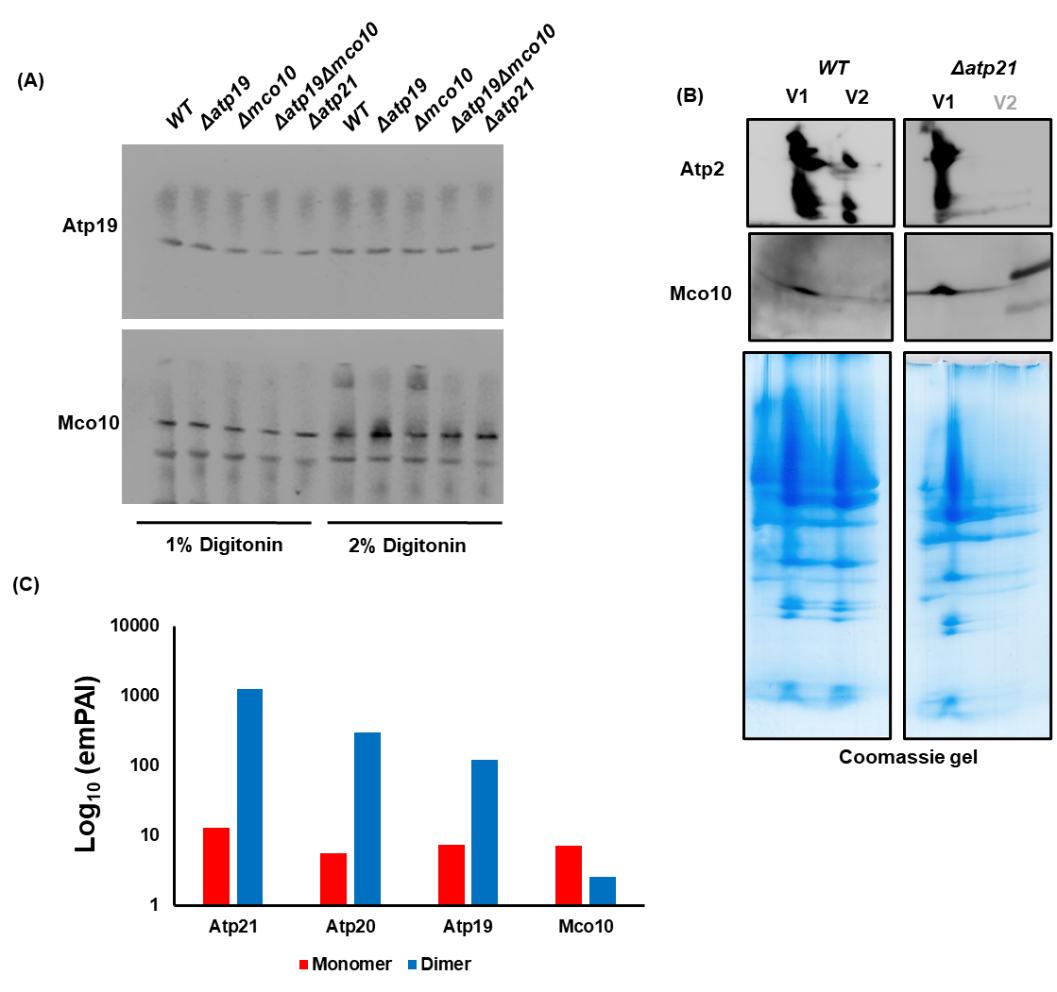

**(A)**

**(B)**

**(C)**

**Fig S5.** **A.** Mco10 and Atp19 antibody nonspecifically bind to monomer or dimer extracted by digitonin and separated in BN PAGE. **B**. Monomer and dimer subunits from the wild type and *Δatp21* from two-dimensional BN-SDS-PAGE gel separation were transferred to PVDF membrane and visualized by western blotting with respective antibodies. Mco10 is also detected in the monomer when Atp21 is deleted. **C.** Protein abundance index (emPAI) values of Atp19/*k*, Atp20/*g*, Atp21/*e* and Mco10 as determined from interactome analysis of the monomers and dimers of ATP synthase in *S. cerevisiae.*

**3**

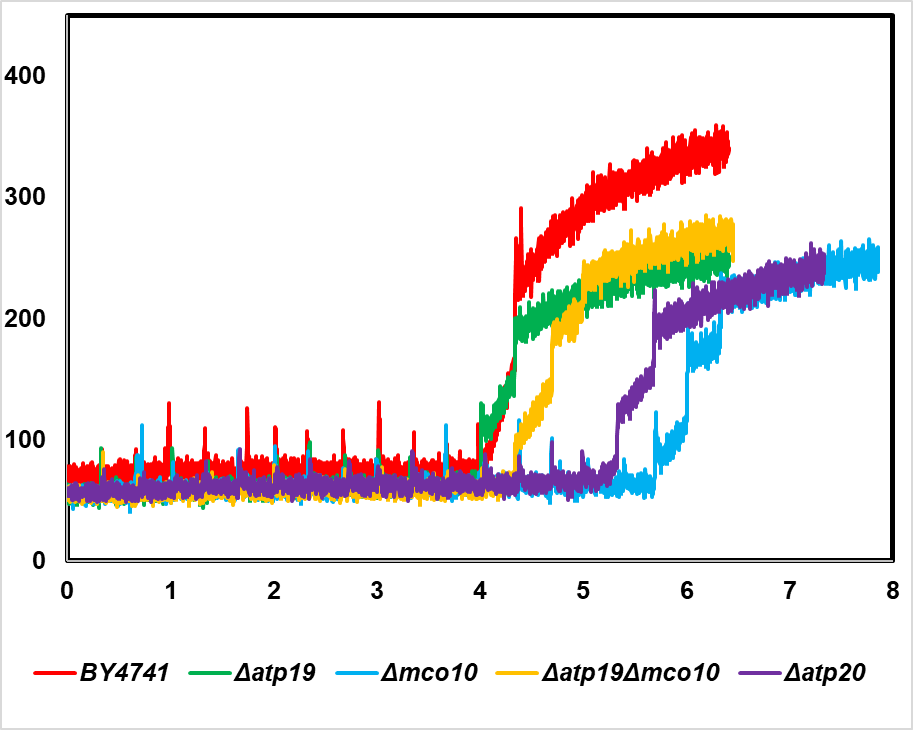

**Fluorescence intensity (A.U.)**

**Time (mins)**

**Fig S6.** The yPTP induction time measured in the CRC experiment. 1 mg of mitochondria was added to CRC buffer containing the calcium ionophore ETH129 and the calcium green-5N calcium indicator, and 10 µM CaCl_2_ was added every 20 secs intervals until the mitochondria stops taking calcium and rapidly releases it into the buffer. Traces are representative of at least 3 independent experiments

**4**
